# Engineering of NEMO as calcium indicators with large dynamics and high sensitivity

**DOI:** 10.1101/2022.08.23.504677

**Authors:** Jia Li, Ziwei Shang, Jia-Hui Chen, Wenjia Gu, Li Yao, Xin Yang, Xiaowen Sun, Liuqing Wang, Tianlu Wang, Siyao Liu, Jiajing Li, Tingting Hou, Dajun Xing, Donald L. Gill, Jiejie Li, Shi-Qiang Wang, Lijuan Hou, Yubin Zhou, Ai-Hui Tang, Xiaohui Zhang, Youjun Wang

**Affiliations:** Beijing Key Laboratory of Gene Resource and Molecular Development, College of Life Sciences, Beijing Normal University, Beijing, 100875, China; State Key Laboratory of Cognitive Neuroscience & Learning, IDG/McGovern Institute for Brain Research, Beijing Normal University, Beijing, 100875, China; Hefei National Research Center for Physical Sciences at the Microscale, CAS Key Laboratory of Brain Function and Disease, and Ministry of Education Key Laboratory for Membrane-less Organelles & Cellular Dynamics, Division of Life Sciences and Medicine, University of Science and Technology of China, Hefei 230026, China; Exercise physiology and Neurobiology Lab, College of P.E. and Sports, Beijing Normal University, Beijing, 100875, China; Institute of Biosciences and Technology, Texas A&M University, Houston, TX, 77030, USA; State Key Laboratory of Membrane Biology, College of Life Sciences, Peking University, Beijing 100871, China; Department of Cellular and Molecular Physiology, Pennsylvania State University College of Medicine, Hershey, Pennsylvania 17033; Department of Translational Medical Sciences, School of Medicine, Texas A&M University, Houston, TX, 77030; Institute of Artificial Intelligence, Hefei Comprehensive National Science Center, Hefei 230026, China; Key Laboratory of Cell Proliferation and Regulation Biology, Ministry of Education, College of Life Sciences, Beijing Normal University, Beijing, 100875, China

## Abstract

Genetically-encoded calcium indicators (GECI) are indispensable tools for real-time monitoring of intracellular calcium signals and cellular activities in living organisms. Current GECIs face the challenge of sub-optimal peak signal-to-baseline-ratio (SBR) with limited resolution for reporting subtle calcium transients. We report herein the development of a suite of calcium sensors, designated NEMO, with fast kinetics and wide dynamic ranges (>100-fold). NEMO indicators report Ca^2+^ transients with peak SBRs ~20-fold larger than the top-of-the-range GCaMP series. NEMO sensors further enable the quantification of absolution calcium concentration with ratiometric or photochromic imaging. Compared to GCaMPs, NEMOs could detect single action potentials in neurons with a peak SBR two times higher and a median peak SBR four times larger *in vivo*, thereby outperforming most existing state-of-the-art GECIs. Given their high sensitivity and resolution to report intracellular Ca^2+^ signals, NEMO sensors may find broad applications in monitoring neuronal activities and other Ca^2+^-modulated physiological processes in both mammals and plants.

---

Genetically-encoded calcium (Ca^2+^) indicators (GECI) are indispensable tools for real-time monitoring of Ca^2+^ signals^1^ and cellular activities ^2^. The signal-to-baseline ratio (SBR; ΔF/F_0_), defined as the ratio of the absolute fluorescence (F) changes (F-F_0_, or ΔF) over the basal fluorescence (F_0_), is a key parameter used to gauge the performance of mono-colored GECIs ^3^. Tremendous efforts have been devoted to generate GECIs with faster kinetics (e.g., jGCaMP8 series ^4^), but progress toward increased maximal fluorescence change has remained relatively lagging since the development of GECO and GCaMP6 series approximately ten years ago ^5,6^.

With a Ca^2+^-sensing module installed within one fluorescent protein (FP), single-FP-based indicators use Ca^2+^-dependent fluorescence changes to report Ca^2+^ transients. Calmodulin (CaM) together with its target peptide (such as RS20 or M13) is among the most commonly-used Ca^2+^ sensing modules. Two strategies have been applied to link CaM-M13 with FP: (i) GCaMP-like design ^6^ to install CaM and M13 to the C- and N-termini of a FP; and (ii) NCaMP7-like strategy ^7^ to insert CaM-M13 into the middle of a FP ^8^. Modifications within the linkers or interaction interfaces among CaM, M13 and FP were proven successful strategies to improve the performance of GCaMP variants ^4,6,9^. Nevertheless, further improvements in dynamic range (DR) are restricted by the brightness of EGFP. While NCaMP7 or mNG-GECO^10^ was built upon the brightest monomeric green FP, mNeonGreen (mNG) ^11^, they exhibited a relatively small *in cellulo* DR ^7^. By combining the advantages of both the GCaMP and NCaMP7 series, we set out to develop substantially improved GECIs with fast speed and high DRs building upon mNG.

## Engineering of m<u>Ne</u>onGreen-based calciu<u>m</u> indicat<u>o</u>rs (NEMO)

Single-FP-based indicators share some structural similarities at the sensing module insertion sites ^812^. Assuming that the design strategies for CaM-based indicators might be transferable in principle among GECIs, we created a series of constructs mostly by applying known GECI design strategies toward mNG, and screened their performance in HEK293 cells (**Fig. 1A–C**, and **Supplementary Tables 1–2**).

**Figure 1.**
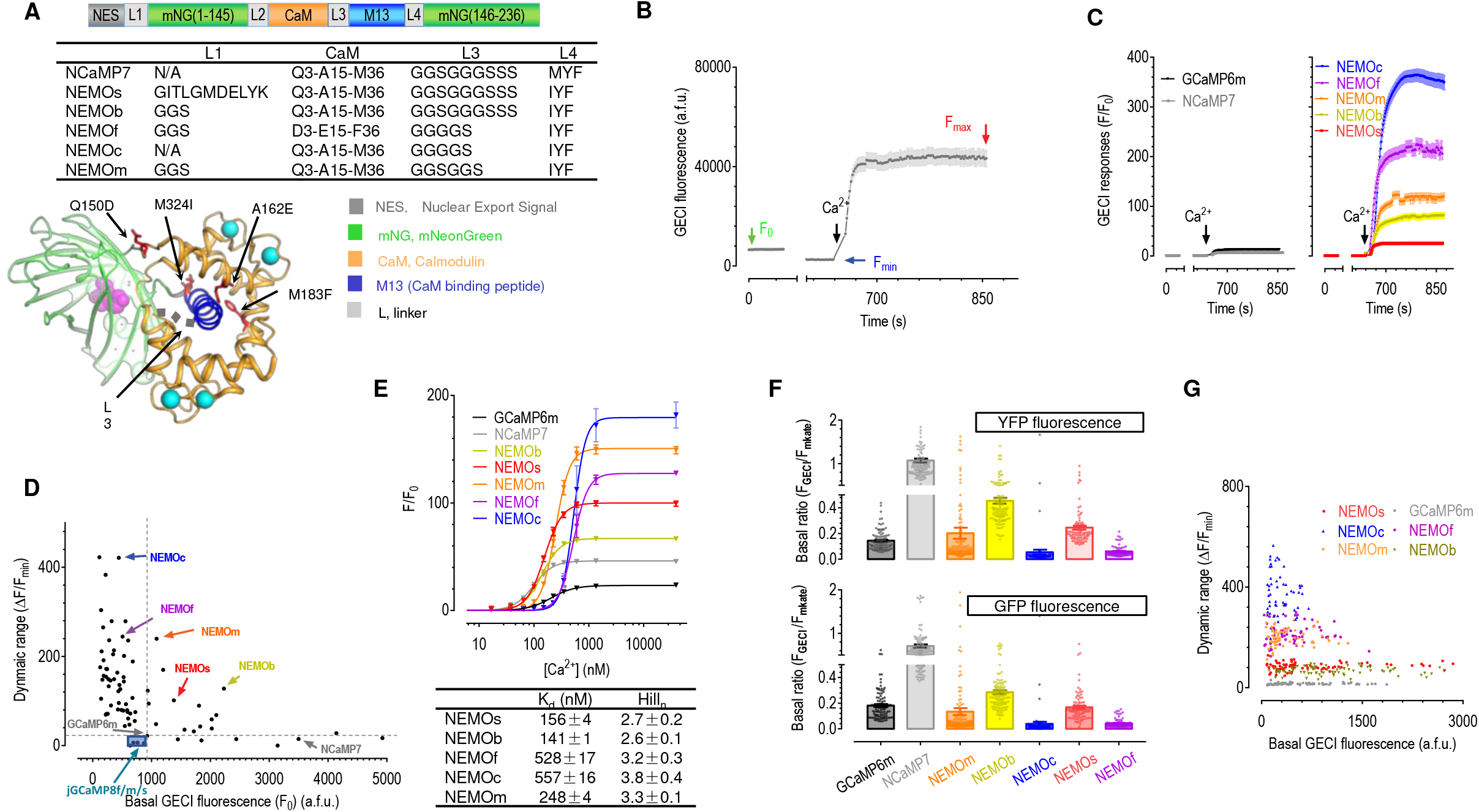
Screening and *in vitro* characterization of NEMO indicators. (A) NEMO sensors are generated by introducing amino acid substitutions in NCaMP7. Top panel, a diagram showing the design of NEMO variants; a table (middle panel) or NCaMP7 structure ^7^ (bottom panel) showing key amino acids substitutions introduced into NCaMP7 to generate NEMO variants. (B–E) Screening of GCaMP and NCaMP7 variants in HEK293 cells. B) Ca^2+^ imaging-based screening. A typical trace from NCaMP7-expressing cells is shown. To avoid saturation of the camera, after recording the basal fluorescence (F0) with regular exposure time (approximately 500 ms), time-series for variants with high dynamic range were recorded using one-tenth to one-fifth the exposure time. Afterwards, the fluorescence response curves of each cell were scaled up according to the corresponding F0. After recording F0, endoplasmic reticulum Ca^2+^ store was depleted using 2.5 μM ionomycin (iono) and 1 μM thapsigargin (TG). Then, the cells were incubated in an imaging solution containing 300 μM EGTA, to read minimal GECI fluorescence (Fmin). Finally, the cells were exposed to imaging solution containing 100 mM Ca^2+^ to obtain the maximal response (Fmax) via store-operated Ca^2+^ entry. C) Representative traces of GCaMP6m and NCaMP7 (left), or selected NEMO sensors (right). D) Scatter plot of F0–mean dynamic range (DR, (Fmax - Fmin) / Fmin) of the indicated GECIs. E) *In vitro* dose–response curves of NEMO sensors. Top, typical traces; Bottom, statistics (see **Supplemental Table 3** for details) (n=3 independent biological replicates; >17 cells per repeat). Data shown as mean ± s.e.m. (F) Basal brightness of NEMO, NCaMP7 or GCaMP sensors viewed with YFP (top) or GFP (bottom) filters. To achieve better estimation of the basal fluorescence of GECIs (FGECI), FGECI of cells expressing mKate-P2A-GECI constructs were normalized against the fluorescence of mKate, an expression marker (FmKate). (GCaMP6m, n=99 cells; NEMOm, n=142 cells; NEMOb, n=119 cells NEMOc, n=89 cells; NEMOs, n=91 cells; NEMOf, n=86 cells; NCaMP7, n=114 cells.) Three independent biological repeats. Data shown as mean ± s.e.m.(G) F0–dynamic range of individual cells expressing NEMO variants or GCaMP6m examined with a YFP filter set.

We first evaluated their basal fluorescence (F_0_) and the ratio between maximal (F_max_) and minimal (F_min_) fluorescence, or DR, (F_max_ - F_min_) / F_min_). To allow measurements of F_min_, the endoplasmic reticulum (ER) Ca^2+^ store was depleted by 10-min incubation with 2.5 μM ionophore ionomycin (iono) and 1 μM thapsigargin (TG, an inhibitor of the sarcoplasmic/endoplasmic reticulum Ca^2+^ ATPase). A high amount of Ca^2+^ (100 mM) was added to the bath to induce F_max_ via store-operated Ca^2+^ entry (SOCE) (**Fig. 1B**), Top candidates (**Fig. 1C, Extended Data Fig. 2**) were identified based on both the F_0_ and DR values (**Fig. 1D**). We found that only the NCaMP7-like ^7^ design (**Fig. 1A**) showed improved dynamics and speed (**Supplementary Table 1&2**). We identified five best-performing constructs and named them as m<u>Ne</u>onGreen-based Calciu<u>m</u> indicat<u>o</u>r (NEMO), including the medium (NEMOm), high contrast (NEMOc), fast (NEMOf), bright (NEMOb) and sensitive (NEMOs) versions (**Fig. 1D, Extended Data Fig. 1 and 23A**).

### *Ex vivo* characterization of NEMO sensors

The overall *in cellulo* DR of NEMO sensors seemed to be superior to that of top-of-the-range GECI proteins tested side-by-side. The DRs of NEMOs or NEMOb (102.3 ± 4.0 or 128.8 ± 3.1, respectively) were found to be at least 4.5-fold higher than that of GCaMP6m or NCaMP7 (**Fig. 1D**). And the DRs of NEMOm and NEMOc were further increased to 240.7 ± 7.6 and 422.2 ± 15.3, respectively, 9.5 to 25.7-fold higher than those of GCaMP6m or NCaMP7 (**Fig. 1D**, and **Supplementary video 1**). To the best of our knowledge, NEMO indicators represent a class of GECI sensors with an exceptional *in cellulo* DR of over 100-fold.

We further assessed the *in vitro* performance of NEMO variants. Except for NEMOb, four NEMO sensors showed DR values either close to (NEMOs) or larger than 100-fold (**Fig. 1E, Extended Data Fig. 2B**). We thus did not focus on NEMOb for further characterization. It is surprising that *in vitro* DR values were smaller than their corresponding in-cell ones, as GCaMP-like design usually show the opposite ^13^. It is likely that macromolecular crowding and reducing condition in cytosolic environment may account for this higher in-cell DR ^14,15^, which warrant follow-on studies in the immediate future.

We next examined the basal fluorescence of NEMO sensors with a P2A-based bicistronic vector to drive the co-expression of mKate (as an expression marker) and GECIs at a near 1:1 ratio. The basal GECI brightness was indicated by the fluorescence ratio of GECI and mKate (**Fig. 1F**). Normalized basal brightness of all NEMO sensors was much lower than that of NCaMP7, and the brightness of NEMOc or NEMOf was only approximately 0.25-0.5 of GCaMP6m. This finding indicates that the lower basal fluorescence of NEMO variants might contribute to the observed large DR of NEMO indicators, in particular for NEMOc and NEMOf. However, even though the DRs of NEMOm, NEMOs and NEMOb were over 5-fold higher than that of GCaMP6m, their basal fluorescence was either similar to or brighter than that of GCaMP6m (**Fig. 1D, 1F**). Hence, high DRs for these three indicators could be attributed to their maximal brightness being larger than that of GCaMP6m as well. In consonance with this notion, NEMO-expressing cells with comparable basal fluorescence to those expressing GCaMP6m still exhibited larger dynamics (**Fig. 1G**). Together, these results establish NEMO indicators as a class of GECIs with extraordinarily large Ca^2+^-dependent changes in fluorescence.

Using NEMOc as an example, we next set out to decipher the mechanisms underlying the superior DR of NEMO sensors (**Extended Data Fig. 3A**). Similar to most GECIs^7 16 17^, the fluorophore of NEMOc adopted two configurations: an anionic state (peak at 509 nm), and a neutral state (403 nm) (**Extended Data Fig. 3B**). The Ca^2+^-induced brightening of NEMOc fluorescence is similarly caused by increasing both the proportion and molecular brightness of anionic form (**Extended Data Fig. 3C–E**, and **Supplementary Table 4**) ^7 16 17^. The increase in the DR was mostly associated with the considerably dimmer anionic fluorophore of NEMOc (0.22 ± 0.01 mM^−1^cm^−1^) in the absence of Ca^2+^, which was approximately one-sixth that of NCaMP7 and one-fifth that of GCaMP6m. Compared to GCaMP6m, the high DR of NEMOc was also a result of increased brightness of Ca^2+^-saturated anionic NEMOc (64.26 ± 2.67 mM^−1^cm^−1^), approximately three times that of GCaMP6m. Moreover, the *in vitro* and *in cellulo* DR of NEMOs under two-photon excitation remained largely similar to those observed with one-photon excitation (**Extended Data Fig. 4A–C**, and **Supplementary Table 5**). Overall, these findings indicate that the superior DR of NEMO indicators can be largely ascribed to the Ca^2+^-dependent fold-of-increase in the molecular brightness of the anionic fluorophores in NEMO variants.

To examine whether it is possible to compensate weaker basal NEMO fluorescence with stronger illumination, we measured the photostability of NEMO sensors. NEMO indicators showed better photostability than GCaMP6m or mNG ^11^ (**Extended Data Fig. 5A**, top left two panels). It endured nearly 40 times (1.52 mW) higher illumination than GCaMP6m (0.04 mW), and showed no apparent photobleaching. The stronger illumination (from 0.04 mW to 1.52 mW) could potentially enhance the basal fluorescence of NEMOm sensor by over 60-fold (**Extended Data Fig. 5A**, top right panel), greatly broadening the applicability of NEMO sensors in scenarios requiring stronger light illumination, such as monitoring Ca^2+^ signals *in vivo* within subcellular compartments with dim NEMOf indicator. Collectively, these results established NEMO variants as photostable biosensors with large DRs.

### Performance of NEMO sensors in non-excitable cells

We next examined the ability of NEMO sensors to report signals induced by submaximal activation of muscarinic acetylcholinergic receptors with carbachol (CCh, 10 µM). As to the CCh-induced Ca^2+^ transients in HEK293 cells, peak signal-to-baseline (SBR, ΔF/F_0_) values of NEMOb and NEMOs were at least three times higher than those of GCaMP6m, jGCaMP8f ^4^, and NCaMP7 (**Fig. 2A, Extended Data Fig. 5B, C**). The SBRs of NEMOm (101.9 ± 6.6), NEMOc (112.0 ± 9.8) and NEMOf (194.3 ± 7.7) were 13-25 times higher than that of GCaMP6m. Similarly, the performance of NEMO sensors in reporting subsequent Ca^2+^ oscillations were also superior over jGCaMP8f and NCaMP7 (**Fig. 2A, Extended Data Fig. 5B-C**, and **Supplementary Video 2**).

**Figure 2.**
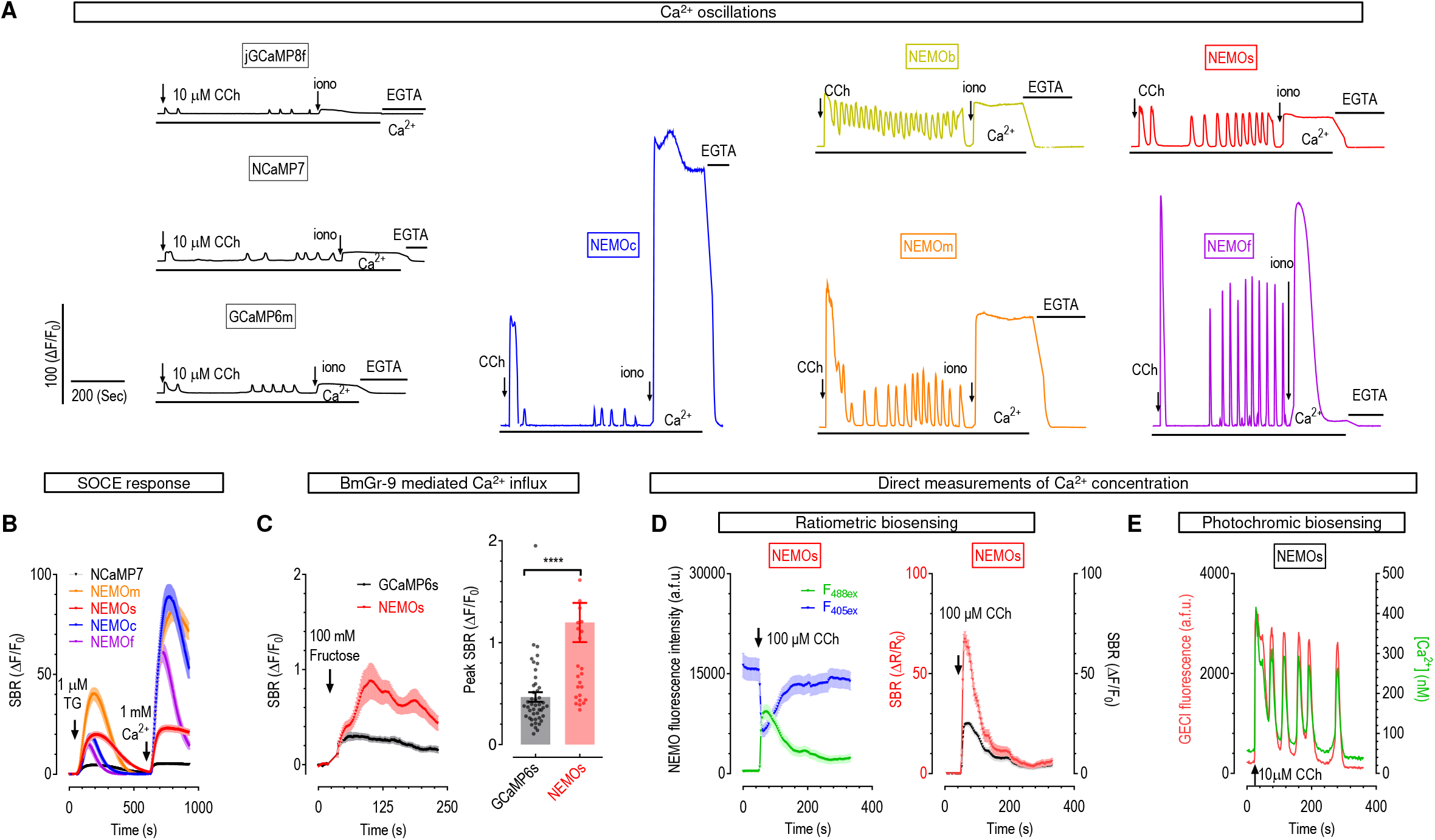
Performance of NEMO sensors in non-excitable mammalian cells. (A) Typical Ca2+ oscillations in HEK293 cells induced by carbachol (CCh, 10 μM), as indicated by GCaMP6m, NCaMP7, and NEMO sensors. n=3 independent biological replicates, with at least 15 cells per repeat. (B) Ca2+ release and store-operated Ca2+ entry (SOCE) responses induced by thapsigargin (TG). n=3 independent biological replicates, with at least 20 cells per repeat. Data shown as mean ± s.e.m. (C) Fructose-elicited response in cells co-expressing BmGr-9, an insect fructose receptor. Left, typical traces; right, statistics (****, p < 0.0001, unpaired Student’s *t*-test, two-tailed) (GCaMP6s, n=45 cells; NEMOs, n=28 cells examined over 3 independent biological replicates.) Data shown as mean ±s.e.m.. (D-E) Measurements of Ca^2+^ concentration with NEMO sensors. D) Ratiometric measurements with NEMOs. NEMOs transients in cells upon excitation at 488 nm or 405 nm, induced by 100 μM CCh. Left, typical NEMOs fluorescence response when excited at 488 nm or 405 nm. Right, representative intensiometric (black) or ratiometric (red, F488ex/F405ex) responses of the same set of cells shown on the left. n = 3 independent biological replicates, with at least 16 cells per repeat. Data shown as mean ± s.e.m.. E) Intermittent photochromism-enabled absolute quantification (iPEAQ) of Ca2+ levels. Shown are fluorescence intensities (purple) and [Ca2+] traces (green) of NEMOs-expressing cells in response to 10 μM CCh. In-cell calibration to determine the absolute Ca2+ concentration, or [Ca2+], was shown in **Extended Data Fig. 4G**. n = 3 independent biological replicates, with at least 9 cells per repeat.

We further examined the performance of NEMO sensors in detecting weak Ca^2+^ signals in HEK 293 cells. In response to TG-induced Ca^2+^ releases and SOCE, the peak SBR values of NEMO sensors were at least 5 times higher than that of NCaMP7 (**Fig. 2B**, and **Extended Data Fig. 5D**). When monitoring Ca^2+^ transients induced by the *Bombyx mori* gustatory receptor (BmGr-9), whose amplitude is much smaller than SOCE ^18^, the responses of NEMO sensors were much stronger than that of GCaMP6s (**Fig. 2C**). Together, intensiometric NEMO responses to the investigated Ca^2+^ signals in non-excitable cells are remarkably larger than those of other GECIs tested side-by-side.

We next examined whether the larger DR of NEMO sensors could enable more sensitive discrimination of Ca^2+^ signals with varying amplitudes by comparing their performance with GECIs bearing comparable Ca^2+^-binding affinities. We first examined GECIs’ responses to the stepwise increase in Ca^2+^ influx induced by a co-expressed optogenetic tool, Opto-CRAC, when subjected to varying photo-activation duration^19–21^. Compared to the GCaMP6m signals, the NEMOm signals were significantly larger, showing a stepwise increase in response to prolonged photo-stimulation (**Extended Data Fig. 6A**, and **Supplementary Video 3**). Second, we compared the graded SOCE signals in responses to increased extracellular Ca^2+^ concentrations. NEMOm or NEMOs could discriminate more external Ca^2+^ gradients than GCaMP6m or NCaMP7 did (**Extended Data Fig. 6B-C**), with SNRs significantly higher than their corresponding counterparts (**Extended Data Fig. 6D-F**).

One major drawback of intensiometric Ca^2+^ sensors is that they could not directly report Ca^2+^ concentration. We thus asked whether NEMO could serve as ratiometric sensors to measure absolute Ca^2+^ concentrations. Indeed, the fluorescence of NEMOs excited by 405-nm light (F_405_) was reduced as a function of increasing Ca^2+^ concentration (**Extended Data Fig. 7A**). Consequently, the *in vitro* DR indicated by the F_490_/F_405_ ratio was 3.4-fold higher than that obtained with F_490_ only (**Extended Data Fig. 7B**). Similarly, the in-cell ratiometric DRs were significantly larger than intensiometric ones (**Fig. 2D**, and **Extended Data Fig. 7C-D**).

Violet light illumination approximately doubled the fluorescence of NEMO sensors under low Ca^2+^ conditions, indicating the existence of a photochromic effect ^22^ (**Extended Data Fig. 7E**). Indeed, further tests showed that brief 405-nm illumination superimposed on 488-nm light could increase NEMOf fluorescence in an inversely Ca^2+^ dependent manner, with the peak fluorescence named as F_0_. After switching off the violet light, NEMOf quickly relaxed back to its basal state, or termed as the minimal fluorescence (F_end_) (**Extended Data Fig. 7F**). One such photochromic cycle would allow the calculation of its photochromism contrast, defined as ((F_0_ - F_end_) / F_0_)_hv_. ^22^. We thus used NEMOs to report Ca^2+^ concentration with the newly-developed intermittent photochromism-enabled absolute quantification (iPEAQ) method ^22^. Using basal photochromism contrast together with two *in vitro* Ca^2+^-titration curves (**Extended Data Fig. 7G**), we quantified CCh-induced Ca^2+^ releases in terms of absolute Ca^2+^ concentration (**Fig. 2E**). Collectively, NEMO sensors can be used as ratiometric or photochromic indicators ^22^ to report absolute Ca^2+^ levels.

### Assessing NEMO sensors in neurons and in planta

We next examined the responses of NEMO sensors in dissociated rat neurons excited by electrical field stimulation with a GCaMP-compatible imaging setup (**Fig. 3**). We observed that all NEMO sensors were able to detect Ca^2+^ signals elicited by a single action potential (AP) (**Fig. 3A**), with peak SBR approximately twice as high as GCaMP6s or GCaMP6f. Consistent with its *in vitro* K_off_ value being higher than that of GCaMP6f (**Supplementary Table 3**), NEMOf was fast enough to discriminate neuronal responses stimulated with a frequency up to 5 Hz (**Fig. 3B**). Thus NEMO sensors perform similarly or better than the existing EGFP-based sensors in terms of single-AP detection sensitivity and response time.

**Figure 3.**
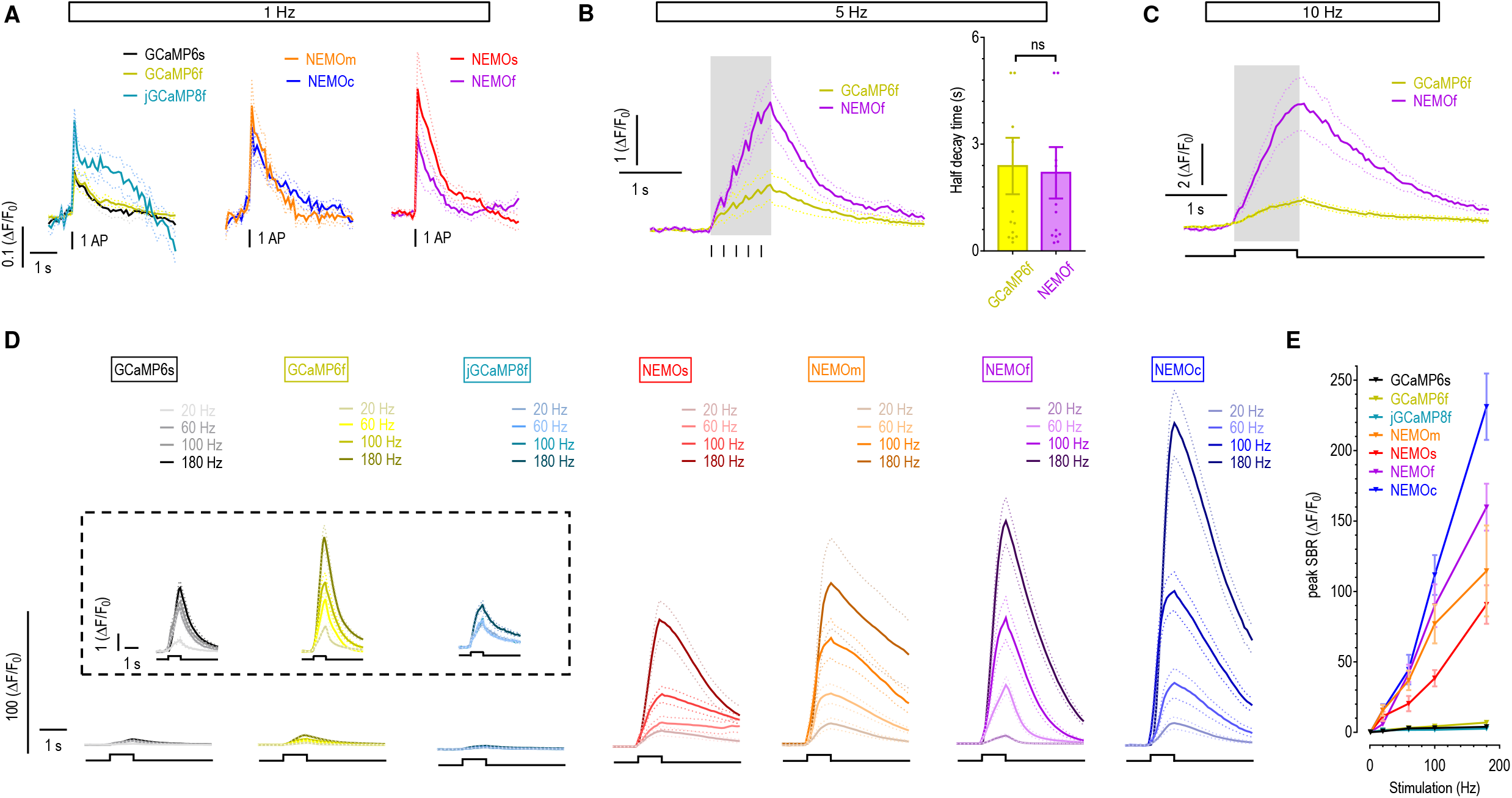
Electric field stimulation-induced NEMO responses in rat hippocampal neurons. (A) Average Ca^2+^ responses reported by GECIs 1 Hz stimulation. (mean traces, n=10-11 cells) (B-C) Mean NEMOf and GCaMP6f transients induced by 5 Hz (B, left, mean traces; right, statistics. p = 0.86, unpaired Student’s t-test, two-tailed, n=12 cells) or 10 Hz (C, mean traces, n =11 cells) stimulation. Data shown as mean ± s.e.m. (B-C) Mean GECI responses elicited by stimulation at varied frequencies. Inset, enlarged views of responses of reference GECI sensors (SBR magnified by 9 times). (E) Statistics for data shown in panels A and D. Each GECI measurement set was analyzed from multiple dendrites of at least 10 neurons in three different primary hippocampal neuron cultures. All data in this figure were shown as mean ± s.e.m.. (For stimulation at varied frequencies, GCaMP6s, n=10,10,10,11,10 cells; GCaMP6f, n=10, 11, 11, 11, 10, 10 cells; jGCaMP8f,n=10, 10, 11, 11, 11 cells; NEMOm, n=10, 12, 12, 13, 12 cells; NEMOs, n=10, 10, 11, 14, 10 cells; NEMOf, n=11, 11, 11, 12, 14 cells; NEMOc, n=11, 11, 11, 13, 12 cells.).

Similar to what we observed in non-excitable cells, the high DR of NEMO sensors enabled high-resolution detection of Ca^2+^ signals of various amplitudes. In response to a 5 Hz field stimulation, the peak amplitude of NEMOf response was approximately three times that of GCaMP6f, placing NEMOf among the most sensitive and fast GECIs that include XCaMP^23^, jGCaMP7^9^, or jGCaMP8 series^4^. As the stimulus frequency increased, the difference between peak NEMOf and GCaMP6f responses became more pronounced (5 or 22.7 times higher than that of GCaMP6f; **Fig. 3C-D**), often with their SNRs significantly larger than their counterparts (**Extended Data Fig. 8A**). The NEMO responses were also fairly linear with no apparent saturation even up to 180 Hz (**Fig. 3E**), indicating that they are suitable for resolving neuronal Ca^2+^ dynamics in response to the full spectrum of activities under physiological conditions.

Combining two-photon imaging and whole-cell electrophysiological recording, we further tested NEMO sensors in cortical neurons in acutely-prepared mouse brain slices. (**Extended Data Fig. 8B**, left panel). Under a whole-cell patch clamp condition, the intracellular environment could be perturbed and the GECI signal was diluted ^23^. Despite of this caveat, the responses of all NEMO sensors induced by AP occurring at 50 Hz or higher frequencies were significantly higher than those of GCaMP6, with NEMOf exhibiting the fastest kinetics (**Extended Data Fig. 8B**). Of note, most fast GECI sensors developed to date ^24, 25, 23, 4^ display rather limited DRs. By contrast, NEMOf expressed in both non-excitable and excitable cells responds fast with high DR, making it an ideal tool to decode fast and highly dynamic Ca^2+^ signals in living cells and tissues.

We also tested the usability of NEMO sensors in detecting subcellular Ca^2+^ signals in the leaves of *Arabidopsis thaliana*. Our super-resolution imaging results showed that NEMOm fused to the plasmodesmata-localized protein 1 could readily report Ca^2+^ oscillations near the plasmodesmata (**Extended Data Fig. 9E, Supplementary Video 4)**, a unique structure between plant cells with a diameter about 30~60 nm ^26^. To the best of our knowledge, this is the first report of plasmodesmata-associated Ca^2+^ signals, and opens avenues to probe the molecular machinery that govern calcium dynamics in this specialized subcellular compartment. This finding firmly establishes the feasibility of applying NEMO sensors *in planta*.

### *In vivo* performance of NEMO sensors in rodent brains

We next tested the *in vivo* performance of NEMO sensors with two-photon laser microscopy (**Fig. 4A**) by measuring GECI responses as readout of neuronal activity evoked by drifting grating stimulus in the primary visual cortex^27^. To ensure direct comparison with GCaMP6 sensors, we used 920 nm light, an excitation wavelength optimized for GCaMP but less ideal for NEMO (980 nm) to excite the GECIs. All indicators reported differential changes of visual stimuli (**Fig. 4B**, and **Extended Data Fig. 8C-D**). NEMOf was the fastest among all NEMO variants, with the half decay time (409 ± 54 ms) comparable to GCaMP6f (482 ± 48 ms) (**Extended Data Fig. 8E**).

**Figure 4.**
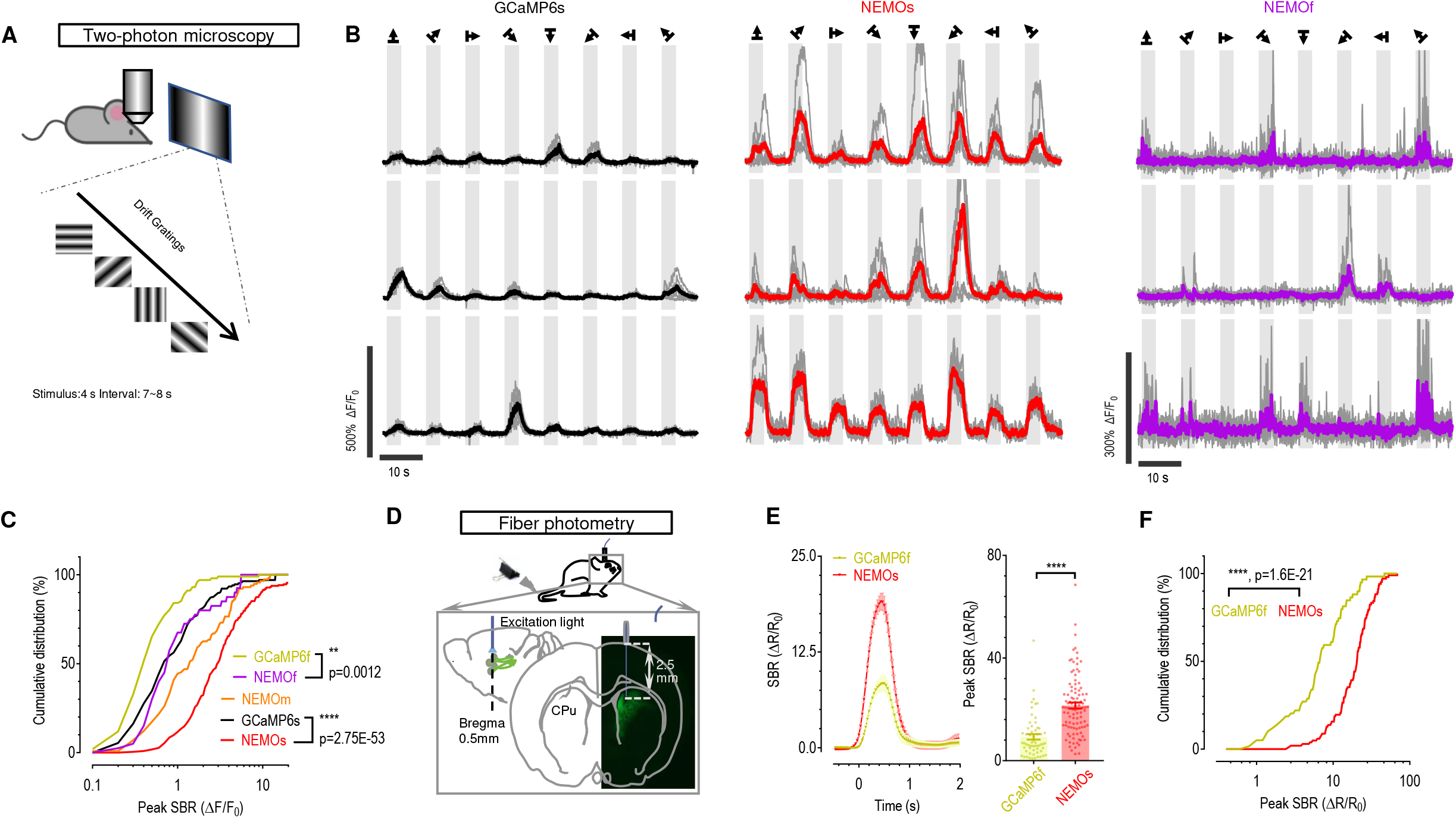
*In vivo* performance of NEMO sensors in monitoring neuronal activities in rodent brain. (A–C) Fluorescence responses in the visual cortex of mice induced by a visual stimulus. A) Diagram showing the experimental setups for two-photon imaging of neurons in response to drift gratings. B) Typical response curves for GCaMP6s, NEMOs and NEMOf. C) Cumulative distribution of peak signal-to-baseline-ratio (SBR) transients of GECI sensors. (For NEMOf versus GCaMP6f and NEMOs versus GCaMP6s, **, p = 0.012; ****, p= 2.75E-53, Kolmogorov–Smirnov test, two-tailed.) (GCaMP6f, n=101 cells from 2 mice; GCaMP6s, n=167 cells from 3 mice; NEMOs, n=223 cells from 4 mice; NEMOm, n=115 cells from 3 mice; NEMOf, n=40 cells from 3 mice.). (D–F) Ratiometric responses of GCaMP6f and NEMOs in neurons of the mouse corpus striatum recorded by fiber photometry. D) Diagram of the experimental setup for fiber photometry recordings. E) Mean ratiometric responses elicited by pinch stimulation at the mouse tail tip. Left, mean traces; right, statistics (****, p < 0.0001, unpaired Student’s *t*-test, two-tailed). NEMOs, data for 102 cells from 6 mice; GCaMP6f, data for 59 cells from 6 mice. Data shown as mean ± s.e.m. F) Cumulative distribution of peak responses shown in E (****, p = 1.6E-21; Kolmogorov–Smirnov test, two-tailed).

We then moved on to compare the sensitivity of GECIs *in vivo*. As to the fraction of responsive cells (**Extended Data Fig. 8F**), no significant difference between NEMO variants and the corresponding GCaMP6 indicators was detected. However, the cumulative distribution of peak ΔF/F_0_ of NEMOm and NEMOs was substantially right-shifted relative to the GCaMP6 signal (**Fig. 4C**), indicating that NEMOm and NEMOs are more responsive. The median response of NEMOs (Δ F/F_0_ =3) was over four and seven times stronger than that of GCaMP6s and GCaMP6f, respectively. The visual-stimuli-induced response reported by NEMOs was much larger than existing values reported by existing sensitive GECIs ^25, 28, 9, 4^. In parallel, the median response of NEMOf (ΔF/F_0_ = 0.80) was significantly larger than that of GCaMP6f (ΔF/F_0_ = 0.44) (right panel in **Fig. 4B** versus **Extended Data Fig. 8C**, left panel), as well as those reported by the known fastest GECIs ^24, 25, 9, 4^.

In addition, NEMOs showed appreciably better SNR (**Extended Data Fig. 8G**) and good basal fluorescence in the mouse V1 that was comparable to GCaMP6s even under excitation conditions optimized for GCaMP (**Extended Data Fig. 9A-B**). NEMOf signal obtained with GCaMP set up showed similar SNR to GCaMP6f (**Extended Data Fig. 8G**). Since the basal NEMOm fluorescence approximately doubled by switching from GCaMP excitation (920 nm) to a NEMO setup (980 nm), NEMOf under optimized illumination (**Extended Data Fig. 9C**) retained its large SBR and showed higher SNR (**Extended Data Fig. 9D**). Since GCaMP6s and NCaMP7 were reported to have similar *in vivo* SNR ^7^, it is likely that optimally-excited NEMOs (i.e., at 980 nm) may exhibit a better SNR than NCaMP7.

Lastly, we recorded NEMOs responses in sensory neurons deeply buried in the mouse brain using fiber photometry and settings optimized for ratiometric GCaMP recordings ^29^. The ratios of GECI fluorescence excited by 410 nm (F_410_) or 470 nm light (F_470_) were used to indicate Ca^2+^ responses of the neurons within the corpus striatum elicited by tail-pinching stimulus (**Fig. 4D**). Even though the near-UV light (410 nm) excitation reduced the dynamics of NEMO and the 470 nm light was not optimal for NEMOs excitation (**Extended Data Fig. 3A** and **Extended Data Fig. 7D**), the median peak response of NEMOs was approximately 3 times that of GCaMP6f (**Fig. 4E**). Collectively, NEMOs and NEMOf are among the most sensitive GECI tools for *in vivo* monitoring of both slow and fast neural activities with a better SNR.

## Conclusions

Here we reported a GECI toolkit with greatly improved photochemical properties. Unlike current indicators that partially sacrifice the dynamic range for improved sensitivity and/or faster kinetics, NEMO variants are fast acting while still retaining superior dynamic ranges to report Ca^2+^ signals. They are more versatile than the most popular GCaMP series, allowing simultaneous imaging with cyan fluorescence, exhibiting higher photostability that can endure substantially stronger illumination, and resisting pH fluctuation better. Overall, the highly sensitive NEMO sensors may serve as the tool-of-choice for monitoring Ca^2+^ dynamics in mammalian cells, tissue, or *in vivo*, as well as *in planta*.

## ACKNOWLEDGMENTS

This work was supported by the National Natural Science Foundation of China (91954205 and 92254301 to Y.W.; 32130043 and 32071025 to X-h. Z.; 31872759 to A.-H. T.; 31971095 to L.H.; and 32171033 to D. X., 91854209, 32230048 to S-Q. W, 92054101 and 32122013 to J. L.), National Science and Technology Innovation 2030 Grants (2022ZD0204900 to X-h. Z.; 2021ZD0202503 to A.-H. T.), the National Key Research and Development Program of China (2020YFA0112200 to A.-H. T.), and the Ministry of Science and Technology of China (2019YFA0802104 to Y.W.), and the China Postdoctoral Science Foundation (2021M703089 to J.C.).

## Author contributions

YW, X-h Z, AT, LH, YZ supervised and coordinated the study. JL designed and generated all the plasmid constructs. JL and WG performed the *in vitro* assays. JL performed all fluorescence imaging in HEK cells, with help from WG and LW. TW and SL performed the optogenetic experiments. JC performed confocal imaging of dissociated neurons. TH performed confocal imaging of dissociated cardiac cells. LY performed two-photon Ca^2+^ imaging of whole-cell patched neurons in mouse brain slices. JL made the transgenetic *Arabidopsis thaliana* and carried out super resolution imaging of plasmodesmata. ZS performed *in vivo* two-photon Ca^2+^ imaging of visual cortical neurons in mice. XS monitored basal fluorescence of indicators in mice visual cortex. XY performed fiber photometry *in vivo*. JL, ZS, JC, WG, LW, LY, ZS, XY and XS analyzed data, with input from the other authors. DLG, SW, DX, and JL provided intellectual input to the manuscript. YZ, X-h Z and YW wrote the manuscript with inputs from all the other authors.

## Conflicts of interests

All authors declare no conflicts of interests.

## Methods

### Plasmids construction

The coding sequence of NCaMP7^7^ was synthesized by BGI Geneland Scientific Co., Ltd, China. The corresponding mutations or substitutions were all included on primers (**Supplementary Table 6**). After PCR, fragments of NEMO were reassembled and inserted into a pCDNA3.1(+) expression vector linearized by BamHI and EcoRI via the Ready-to-Use Seamless Cloning Kit (B632219, Sangon biotech, China). All plasmids were confirmed by sequencing.

### Bacterial expression and protein purification

Transetta (DE3) bacteria (Transgene, Beijing, China) transformed with pET28a plasmids containing coding sequences of sensors were cultured with 300 mM IPTG for 12 h at 20 °C. Pelleted bacterial cells were then suspended in 20 mL buffer 1 (in mM, 20 Tris, 300 NaCl and 1 imidazole, pH 7.2), and subsequently sonicated. Recombinant proteins were then purified by 1 mL Ni Sepharose (17-5318-01, GE Healthcare, USA). Columns were sequentially washed with 20 mL buffer 1 and 10 mL buffer 2 (in mM, 20 Tris, 500 NaCl and 10 imidazole, pH 7.2). Purified proteins were then eluted in 5 mL buffer 3 (in mM, 20 Tris, 100 NaCl and 300 imidazole, pH 7.2) ^9^.

### In vitro Ca^2+^ titrations and kinetic measurements

Ca^2+^ titration: 50 μg/mL GECIs in buffer A (100 mM KCl, 30 mM HEPES, pH 7.2) supplemented with either 10 mM EGTA or 10 mM Ca-EGTA were mixed in various ratios^9^. GCaMP (excitation: 485 ± 5 nm; emission: 510 ± 5 nm) and NEMO (excitation: 490 ± 5 nm; emission: 520 ± 5 nm) fluorescence were measured with a Flexstation 3 microplate reader (Molecular Devices, USA) controlled by SoftMax Pro v7.x. The Ca^2+^ titrations curves were fit with Prism 7 using specific binding with hill slope function.

For measurements of k_off_^9^, 50 μg/mL GECIs in buffer A containing either 10 µM free Ca^2+^ or 10 mM EGTA were rapidly mixed with 1:1 ratio GCaMP and NEMO (excitation 485 nm, emission 520 nm) fluorescence were measured by POLARstar Omega microplate reader (BMG LABTECH, Germany). k_off_ values were calculated in Prism 7 using single exponential regression.

### Measurement of Spectra and quantum yields (Φ) of GECIs

Emission and excitation spectra of purified GECIs in buffer A were recorded with a FS5 spectrophotometer (Edinburgh Instrument, Scotland) controlled by Fluoracle. Absorption spectra were recorded with UV2600 spectrophotometer (Shimadzu, Japan) controlled by UVprobe. Φ was determined with FS5 using 1 cm quartz cuvette. The fluorescence spectra of GECIs with different concentrations were recorded to calculate the corresponding total integrated fluorescence intensities (TIF). Linear regression of TIF - absorbance curves were used to derive the slopes (S) of GECIs. Φ was then calculated as: Φ_protein_ = Φ_standard_ × (S_protein_/S_standard_)^5^. For anionic chromophore 470 nm excitation; reference, fluorescein (fluo) in 0.1 M NaOH (φ=0.925)^30^. For neutral chromophore, 405 nm excitation, TOLLES (φ=0.79).^31^.

### Chromophore extinction coefficients (ε)

The absorption spectra of 200 μg/ml GECIs in buffers (in mM, 30 trisodium citrate, 30 borate, with either 10 EGTA or 10 CaCl_2_, with or without 1 MgCl_2_) with varying pH were measured with Flexstation 3. Then the ε values were obtained with the following calculations^16,17^. Briefly, the corresponding absorbance (OD) in protonated (OD_N_, peak absorption at ~400 nm) and deprotonated (OD_A_, peak absorption at ~500nm) states were first obtained from the absorption spectrum curves. Next, the slope S was calculated via linear regression using OD_N_ - OD_A_ values at pH = 7, 7.2, 8, and 9

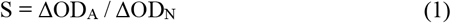

The relationship between the concentration of a chromophore (n), its OD andεis:

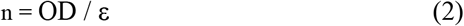

For alkaline-denatured (_D_), protonated (_N_) and deprotonated (_A_) chromophores, the corresponding equation would be n_D_ = OD_D_ / ε_D_, n_A_ = OD_A_ / ε_A_ and n_N_ = OD_N_ / ε_N_. And green GECIs will be totally denatured at pH 12.5, thus n= n_D_. Also n will not change during H^+^ titration, hence

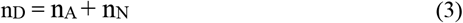

and

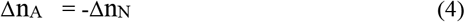

Putting equation (2) or its variants into (3) and (4) would make:

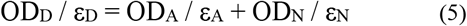

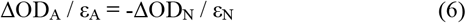

ε_D_ of denatured GFP-like chromophores (44,000 M-1 cm-1) was taken as ε_D_ of green GECIs at pH 12.5 ^1^.

And equation (6) can be rearranged as:

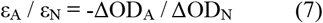

Combining equation (1) and (7), we will obtain

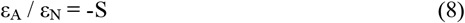

By solving equations (5) and (8), we can get

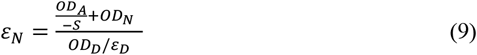

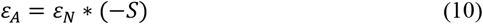

And

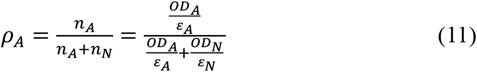

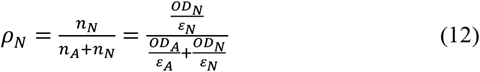

The pKa values were calculated with the ‘specific-binding-with-Hill-slope’ function using Prism 7.

### Two-photon imaging

This was undertaken with a ZEISS LSM 880 NLO microscope equipped with 40x water immersion objective (NA 1.0), gallium arsenide phosphide photomultiplier tubes and ZEN 2.1 software.

*In vitro* excitation spectra: Two-photon cross-section (δ) values were determined ^32^ using the same imaging solutions as those in Φ measurements.

The specific calculation steps are as follows^23^. The time-averaged fluorescence photon flux < F(t) > is given as:

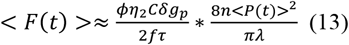

where δ of GECIs can be calculated as:

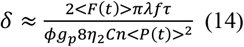

Φ is the fluorescence collection efficiency of the measurement system. ŋ represents the quantum yield, ŋ_2_ is obtained by two-photon excitation and ŋ_2_ derived from one-photon excitation (we set ŋ_1_ = ŋ_2_ = 0.925 for fluorescein, assuming that ŋ was independent of excitation wavelengths. For ŋ_1_ of GECIs, please see Table S4). C is the concentration of the sample. δ is the two-photon cross-section. g_p_ is the degree of second-order temporal coherence, which depends on the pulse shape. f is the pulse repetition rate. τ is the temporal pulse width. n is the refractive index of the sample medium (n =1.3334 for water). < P(t) > is the time-averaged instantaneous incident power of the laser beam (we consider this as a constant), and λ stands for the excitation wavelength.

As 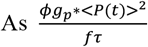 value is a system-specific constant at a certain wavelength setting, we could obtain δ_GECI_ (δ of GECIs) by using fluorescein at 750-990 nm as a standard^33^. According to the above method, we measured the < F(t) > values of each GECI and calculated the δ_GECI_ values of them by equation 15:

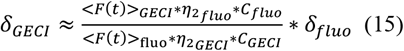

Time-lapse Ca^2+^ imaging of HEK293 cells was acquired using 910 nm (for GCaMP6m) or 970 nm (for NCaMP7 or NEMOs) excitation, and the resulting emission between 500 nm and 600 nm was collected every 2.5 s.

### Cell culture and transfection

HEK293 and HeLa cells (ATCC, cat#: CRL-1573 and CL-0101, respectively) were cultured regularly in DMEM supplemented with 10% FBS (and 5% penicillin and streptomycinat 37°C with 5% CO_2_ ^34,35^. Transfection was performed by electroporation using the Bio-Rad Gene Pulser Xcell system. Transfected cells were seeded on round coverslips and cultured in OPTI-MEM medium containing 7% FBS. All the experiments were carried out 24 h after transfection.

Hippocampal tissue from E18 Wistar rats was first digested with 0.1% trypsin at 37°C for 15 min, washed with mixed complete medium (L-Glutamine DMEM-F12 + 10%FBS) and then pipetted with a Pasteur glass tube (15 mm) to get cell suspension. Neurons were then plated at 50k cells/18 mm coverslips coated with 0.1% poly-D-lysine and cultured in a 37°C, 5% CO_2_ incubator. Half of the media was replaced with neurobasal medium (containing 2% B27 supplement, 0.5mM GlutaMAX-100X) without serum 24 h later, and once per week thereafter. Arac (10 µM) was added 7 days after plating to inhibit gliacyte. Neurons were transfected with GCaMP6 or NEMO-encoding plasmids at 7 to 9 days after plating using a calcium phosphate transfection kit (Takara Bio Inc).

### Fluorescence imaging

Time-lapse fluorescence imaging experiments were carried out using a ZEISS observer Z1 imaging system controlled by SlideBook 6.0.23 ^12^. Cells were imaged in Ca^2+^ imaging buffer (in mM, 107 NaCl, 7.2 KCl, 1.2 MgCl_2_, 11.5 glucose, and 20 HEPES-NaOH, pH 7.2) every 1 or 2 s. Filters: NEMO/NCaMP7, 500 ± 12 nm _ex_/542 ±13 nm _em_; GCaMP, 470 ± 11 nm _ex_/512 ±13 nm _em_..

### Confocal imaging and iPEAQ measurements

Confocal imaging was undertaken with a ZEISS LSM880 microscope equipped with a 63x oil objective (NA 1.4) and ZEN 2.1 software. Excitation was set at 488 and 405 nm, with the emission collected at 500 nm ~ 580 nm.

For the intermittent photochromism-enabled absolute quantification (iPEAQ) biosensing^36^, an *in vitro* Ca^2+^ titration curve was first generated. NEMOs fluorescence in solutions with different free Ca^2+^ concentrations were imaged with the same as those for live-cell imaging. 8 photochromic cycles induced by 2.5 s illumination with 405 nm light were superimposed on 488 nm light to obtain parameters needed to calculate photochromism contrast. The photochromism contrast is defined by peak fluorescence (F_0_) in the presence of 405 nm light together with 488 nm light, and minimal fluorescence (F_end_) in the presence of 488 nm light only: photochromism contrast =((F_0_ - F_end_) / F_0_)_hv_, or (ΔF / F_0_)_hv_. Afterwards, dose-response curves (shown as (ΔF / F_0_)_hv_) and NEMOf fluorescence were plotted and fit with a Hill equation.

In-cell iPEAQ recording of NEMOs were similarly imaged with 488 nm laser. First, 5 photochromic cycles with 2.5 s illumination at 405 nm were superimposed on 488 nm light to obtain basal (ΔF/F_0_)_hv_. Then these values of each single cell were applied to the (ΔF/F_0_)_hv_ Ca^2+^ titration curve to obtain basal [Ca^2+^]. Afterwards, the measured basal NEMOf fluorescence and calculated basal [Ca^2+^] were applied to the NEMOf fluorescence Ca^2+^ titration curve to obtain the normalizing factor needed to convert in cellulo NEMOf fluorescence to the corresponding *in vitro* NEMOf fluorescence, and then to real-time [Ca^2+^]^36^.

### Light-tunable activation of Ca^2+^ entry in HeLa cells

HeLa cells co-transfected with Opto-CRAC (mCherry-LOV2_404-546_-STIM1_336-486_) and NEMOm or GCaMP6m were plated on glass-bottomed dishes (#D35-20-0-TOP, Cellvis), imaged using a Nikon Ti2 Inverted microscope with a Yokogawa W-1 dual spinning disk scanhead, Micro-Scanner for photo-stimulation and stage top incubator for live-cell imaging^19^. To prevent pre-activation of Opto-CRAC, we acquired Ca^2+^ signals (NEMOm or GCaMP6m) under emission light of 525 nm with 1% excitation strength of 488 nm and 10 msec exposure time. To tune the activation of Opto-CRAC, Nikon “A1 Stimulation” toolbar was applied with 488 nm stimulation (2% strength). Varied exposure times were applied in “A1 Stimulation” to control Opto-CRAC activation and photo-induced Ca^2+^ influx.

### Ca^2+^ imaging in neurons

17-20 DIV neurons were imaged using a W1 spinning disc confocal microscope (Ti2-LAPP-Ti2, NIKON) with a 100X oil objective (1.45NA, NIKON) in Tyrode’s solution (in mM:129 NaCl, 5 KCl, 30 glucose, 25 HEPES-NaOH, pH 7.4, 1 MgCl_2_ and 2 CaCl_2_). Field stimulations were performed in a stimulation chamber (Warner Instruments, RC-49MFSH) with a programmable stimulator (Master-8, AMPI). Samples were excited with 488 nm laser and fluorescence was collected with a Zyla4.2 sCMOS camera (Andor) by NIS-Elements AR 5.10.00 software.

### Data analysis for Ca^2+^ imaging in cells or cultured neurons

The corresponding mean fluorescence of regions of interest were analyzed by Matlab 2014a (The MathWorks, Natick, MA, USA) and plotted with Prism 7 software (The Matlab codes are available upon request).

### Ca^2+^ imaging and electrophysiology in cortical slices

Cortical slices were prepared from the adult mice (> P50, either of sex) three weeks after the stereotaxic injection of various AAV vectors encoding different Ca^2+^ probers to the primary visual cortex (V1), following a protocol described in our previous studies^37,38^. Ca^2+^ probe–expressing pyramidal cells (PCs) in cortical slices were recorded in a two-photon laser scanning microscopy system (model FV1200MPE, Olympus) equipped with a wavelength-tunable Mai-Tai femtosecond infra-red laser (DeepSee, Spectra Physics). Whole-cell current-clamp recording on individual PCs was made with the glass micropipettes, filled with the internal solution containing (in mM): 130 K-gluconate, 20 KCl, 10 HEPES, 4 Mg_2_ATP, 0.3 Na_2_GTP, and 10 Na_2_-phosphocreatine (pH= 7.25-7.35, adjusted with KOH, 305±5 mOsm), and a microelectrode amplifier (Axon MultiClamp 700B, Molecular Devices). Membrane potentials were low-pass filtered at 10 kHz, digitized at 20 kHz (DigiData 1440A, Molecular Devices) and acquired by the pClampex 10.3. Simultaneously time-lapse imaging of Ca^2+^ fluorescence signals in the soma of recorded PCs, evoked by action potentials at various frequencies or numbers, was acquired by 17-20 Hz at a 256 × 256 pixel resolution using a 40X water-immersion objective (LUMPlanFL, NA 0.8, Olympus) and the 920 nm laser wavelength. Ca^2+^ signals for each tested action-potential (AP) trains were averaged by 3 sweeps. The signals of neuronal action potential and Ca^2+^ fluorescence were synchronized with an analog connection unit (FV10-ANALOG, Olympus).

The acquired time-lapse images were analyzed offline with the OLYMPUS FV10-ASW 4.2 (Olympus) and our custom MATLAB (MathWorks) scripts. A subtraction of the background fluorescence region outside the PC soma was made to estimate the basal fluorescence intensity *F*_*0*_, and the average *F*_*0*_ for a 0.5-s duration before the onset of action potential trains was used in the calculation of *ΔF/F*_*0*_^39^. The latter quantification process was the same as that used in a previous study^9^.

### In vivo two-photon laser Ca^2+^ imaging of mice V1 neurons

#### Animal Surgery and Virus injection

The use of animals was approved by the Institutional Animal Care and Use Committees (IACUC) of Beijing Normal University, Peking University and University of Science and Technology of China. Mice at postnatal 15-20 days (P15-P20) were transcranially injected with 700 nl *AAV-Syn-GCaMP6f, AAV-Syn-GCaMP6s* or *AAV-Syn-NEMO* virus in the V1. After 3 weeks, animals were used to perform *in vivo* and *ex vivo* calcium imaging experiments. A small craniotomy (2.5 mm ×2.5 mm) was made on the V1 area (centered 2.5 mm left, and 0.5 mm anterior to lambda suture) in the mouse under anesthesia ketamine/ medetomidine (50 mg/kg, 0.6 mg/kg; i.p.)^40^ and then covered with a 3-mm diameter round glass-coverslip. A chamber around the craniotomy, made by dental cement, was filled with the Ringer’s solution containing (in mM): 123 NaCl, 1.5 CaCl_2_, 5 KCl. After a 30-min recovery from the surgery, mice were transported to the two-photon-laser imaging setup.

#### Visual stimuli

Visual stimuli were generated by a custom-developed software using LabVIEW 8.5 (National Instruments, USA) and MATALB (Mathworks, USA), and were presented on a liquid crystal display (LCD) monitor (ThinkVision, Lenovo, China) for in vivo calcium imaging experiments^40^. The simulation covered 0°-80° horizontal visual filed and −35°-40° vertical visual field. Full screen drifting gratings with 8 different orientations (spatial frequency, 0.02 Hz/degree; temporal frequency, 2 Hz, 100% contrast) were presented in a pseudo-random order, and each orientation with 4-s duration was for 6 times at intervals of 7-8 s blank gray-screen stimulus (with identical mean luminance).

#### Two-photon imaging and Ca^2+^ signal analysis

Time-lapse Ca^2+^ imaging from the V1 layer 2/3 PCs was conducted with Scanbox 4.1 system, or a custom-modified Olympus two-photon laser scanning microscopy system (model FV1200MPE). The excitation wavelength was set at 920 nm. Fluorescence signals were acquired at 11-14 Hz. For each acquired fields of view (FOV), regions of interest (ROIs) were manually set on visually-identified neuronal cell bodies.

Ca^2+^ signal analysis was performed, using our custom MATLAB scripts ^9^. F is instantaneous fluorescence, the averaged baseline fluorescence intensity of 1-s duration before visual stimulation onset was calculated as F_0_, and Ca^2+^ responses were defined as ΔF/F_0_=(F-F_0_)/ F_0_. GECI responses evoked by optimal visual orientation stimulus with *p* values < 0.01 (student t-test) were identified as responsive cells. For each responsive cell, the peak ΔF/ F_0_ responses, the peak signal to noise ratio (SNR), and the half-decay time of the maximal ΔF/ F_0_ responses were calculated, respectively, as follows.

For the half-decay time, an exponential function was used to fit ΔF/F_0_ responses of GCaMP6s, NEMOm or NEMOs that were averaged over six trials with optimal stimulus, while the same function was used for maximal peak ΔF/F_0_ responses of GCaMP6f or NEMOf from one trial of optimal stimulus.

The peak SNR was calculated as

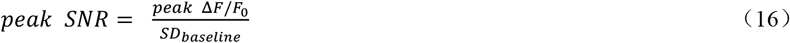

Where *SD*_*baseline*_ is the standard deviations of *ΔF/F*_*0*_ responses before 1-s visual stimulus presentation.

The Orientation Selectivity Index (OSI) and Direction Selectivity Index (DSI) were calculated by using mean *ΔF/F*_*0*_ amplitude (averaging the top 25% of *ΔF/F*_*0*_ responses during 4-s stimulus presentation) over 6 trials evoked by individual 8 orientation stimulation. The OSI was calculated as:

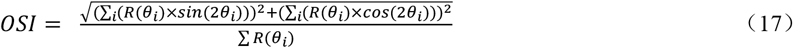

where *θ*_*i*_ is orientation of drifting gratings, (*θ*_*i*_) is the mean *ΔF/F*_*0*_ amplitude at *θ*_*i*_.

The DSI was calculated as

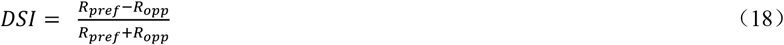

Where *R*_*opp*_ is the mean *ΔF/F*_*0*_ amplitude at the opposite angle to the preferred angle.

### Tail-pinching stimulus and optical fiber recording

After 3 days of adaptive feeding, six male C57BL/6 mice (7 weeks old, 20-25 g) were randomly divided into GCaMP6f group and NEMOs group (N=3). *AAV-Syn-GCaMP6f* or *AAV-Syn-NEMOs* virus were injected into the striatal region (AP: +0.5 mm, R: 1.8 mm, DV: −2.5 mm), respectively. Two weeks after transfection, the experiment was performed using the Reward R810 dual-color multi-channel optical fiber recording system controlled by ORFS V2_14397.

Pinch stimulation at the tail tip was given by a 15-mm long tail clip. 410/470 nm excitation light (17.5/65 μw) and emission between 500-550 nm were transferred via an optical fiber implanted into the virus injection area of anesthetized mice fixed on a stereotaxic instrument. GECI signal excited with 410 nm was a Ca^2+^-independent reference to cancel out motion artifacts^41^.. Pinch-induced GECI responses were recorded by with a rate of 60 fps. One stimulation was defined as one event. For GCaMP6f, n=97; for NEMOs, n=101.

### Animals

All mice were housed in a 12 h light/dark cycle. Food and water were provided ad libitum. The temperature of the room was controlled at 20–25 °C, and the humidity was maintained at 45–60%.

### Super-resolution imaging of plasmodesmata

Full-length cDNA of plasmodesmata-localized protein 1 (PDLP1) was tagged with NEMOm and cloned into pCAMBIA1390. PDLP1 is a well-established plasmodesmata marker^26^. To obtain transgenic plants, Agrobacterium GV3101 containing the resulted construct was transformed into wild type *Arabidopsis thaliana* using the floral dip method^42^. Images were collected from the abaxial leaves of 10-days-old seedlings at 2-s intervals using an Airyscan LSM 880 confocal microscope.

## Data availability

Key NEMO plasmids are available via Addgene (189930 ~ 189934).

All data generated or analyzed during this study are included in this published article (and its supplementary information files).

## Extended Data Figure Legends

**Extended Data Figure 1.**
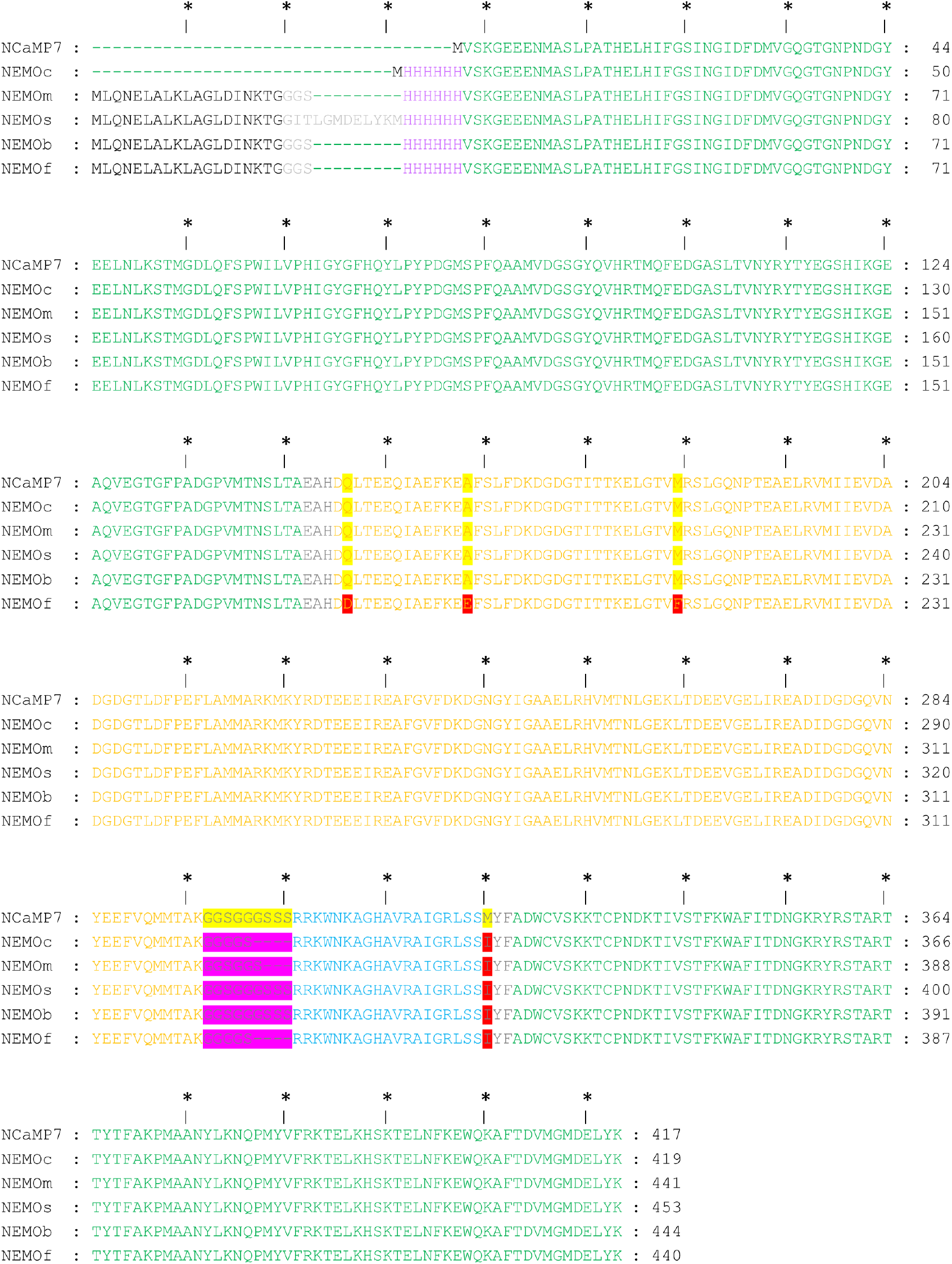
Alignment of the primary sequences of the indicated NCaMP7 and NEMO variants. Mutations in NEMO related to NCaMP7 are highlighted in red. Changes of linker between CaM and M13 of NEMO related to NCaMP7 are shown in pink. The font colors correspond to the colors in the pattern diagram in Figure 1A.

**Extended Data Figure 2.**
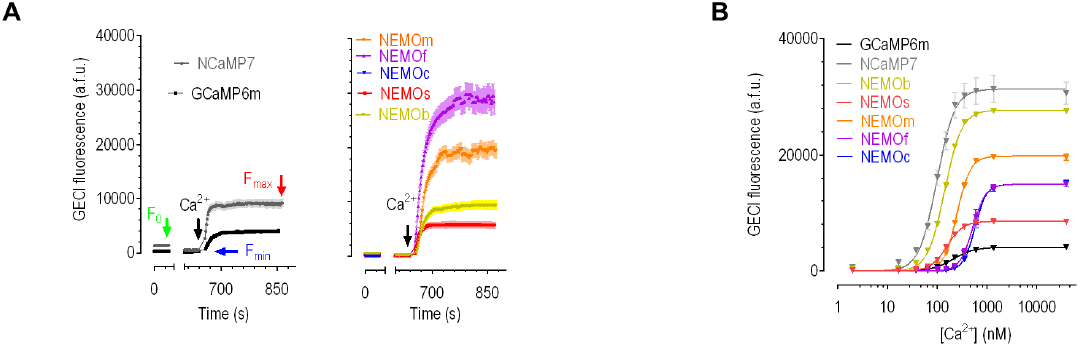
*In cellulo* and *in vitro* responses of NEMO sensors. (A) Typical raw fluorescence readings obtained during the screening of NEMO indicators, corresponding to traces shown in Fig. 1C. GCaMP6m and NCaMP7 (left), or selected NEMO variants (right). (B) Representative raw traces of *in vitro* dose– response curves of GECIs, corresponding to those shown in Fig. 1E. (n=3 independent biological replicates; >17 cells per repeat). Data shown as mean ± s.e.m..

**Extended Data Figure 3.**
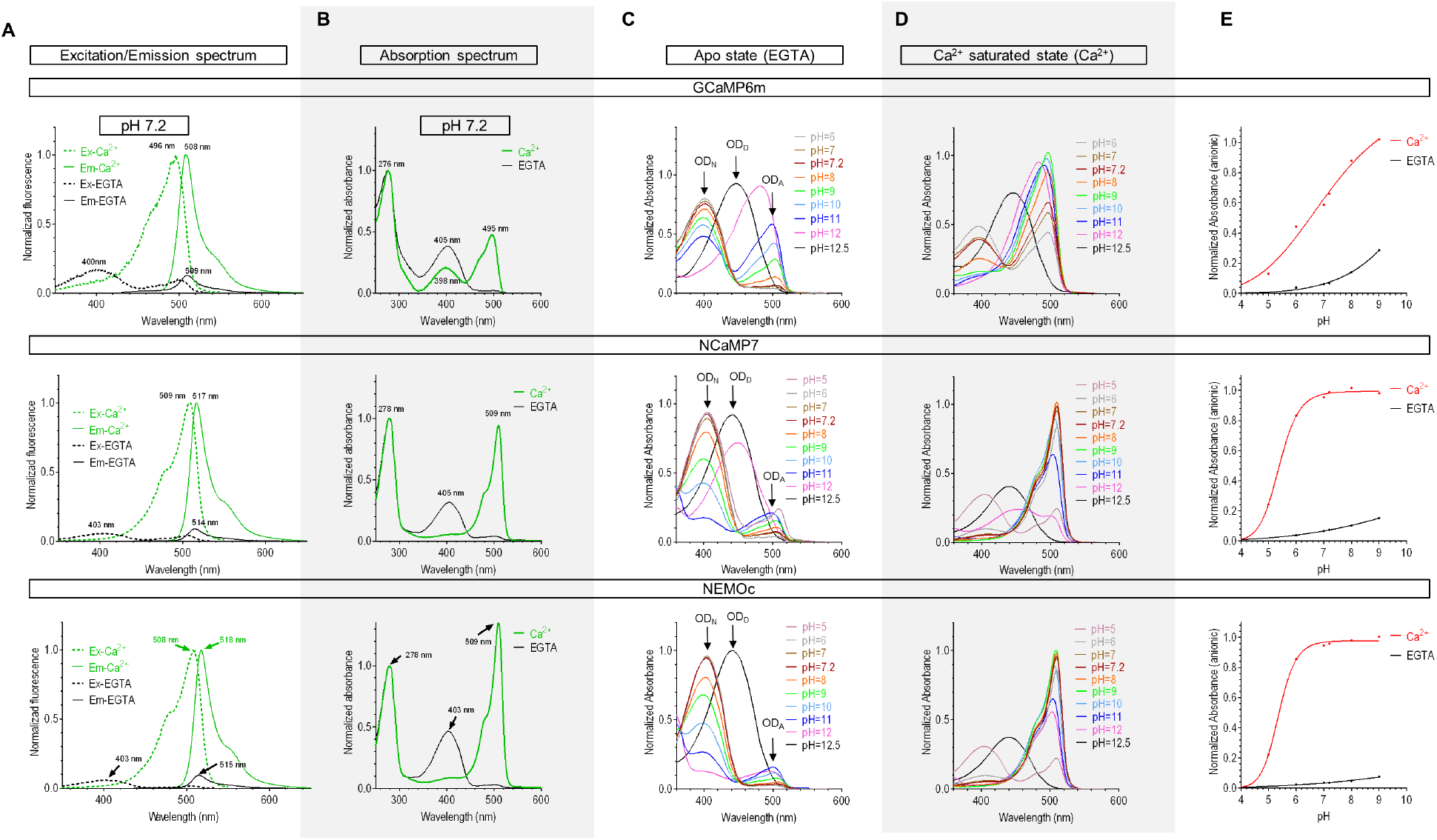
Spectral properties of NEMOc, NCaMP7 and GCaMP6m. (A-B) At pH7.2, typical traces of excitation spectrum, emission spectrum (A), and absorption spectrum (B). (C-D) pH-dependence of absorption spectrum at the apo state (C) and Ca^2+^ saturated conditions (D). (E) Statistics for data shown in panels C&D. n=3 independent biological replicates. The corresponding absorbance (OD) in protonated (OD_N_), deprotonated (OD_A_), and denatured (OD_D_) states were indicated by arrows. n=3 independent replicates.

**Extended Data Figure 4.**
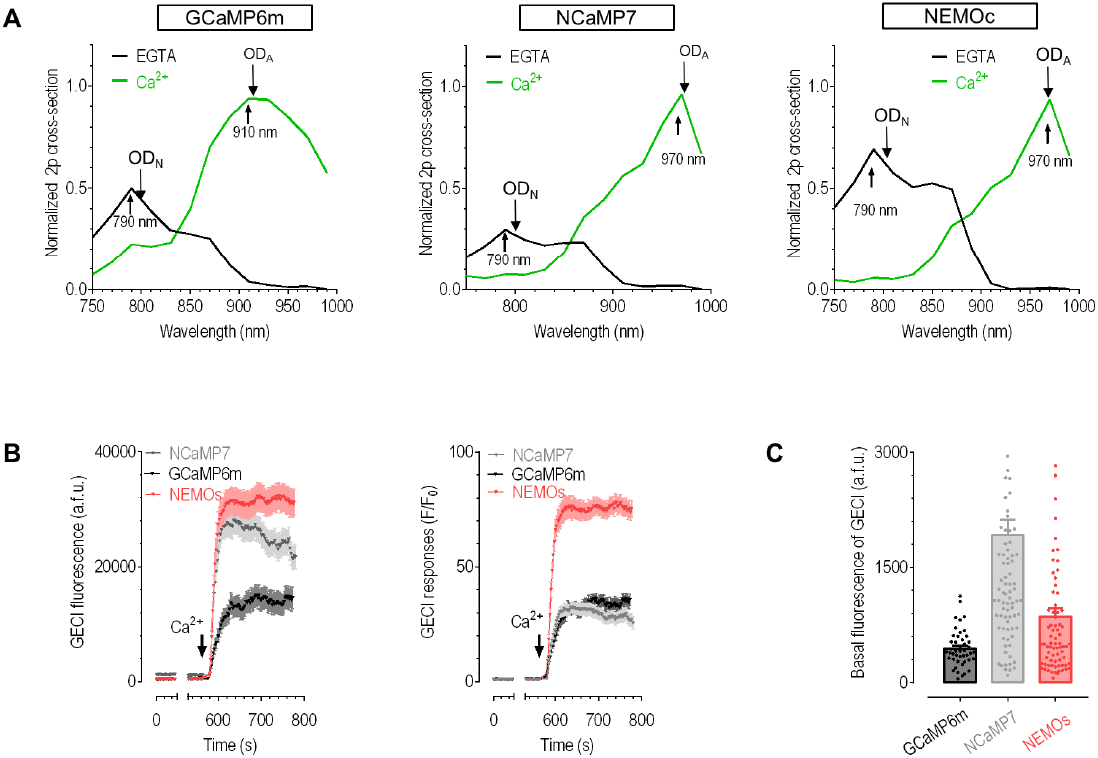
Two-photon spectral properties and performance of NCaMP7, GCaMP6m and NEMO sensors. (A) Typical traces of normalized two-photon action cross-sections at pH 7.2. (B) Dynamic ranges of the indicated GECIs when expressed in HEK293 cells. (C) Statistics of basal fluorescence shown in panel (B). GCaMP6m, n=45 cells; NEMOs, n=76 cells; NCaMP7, n=89 cells examined over 3 independent biological repeats. Data in (B) and (C) were shown as mean ±s.e.m..

**Extended Data Figure 5.**
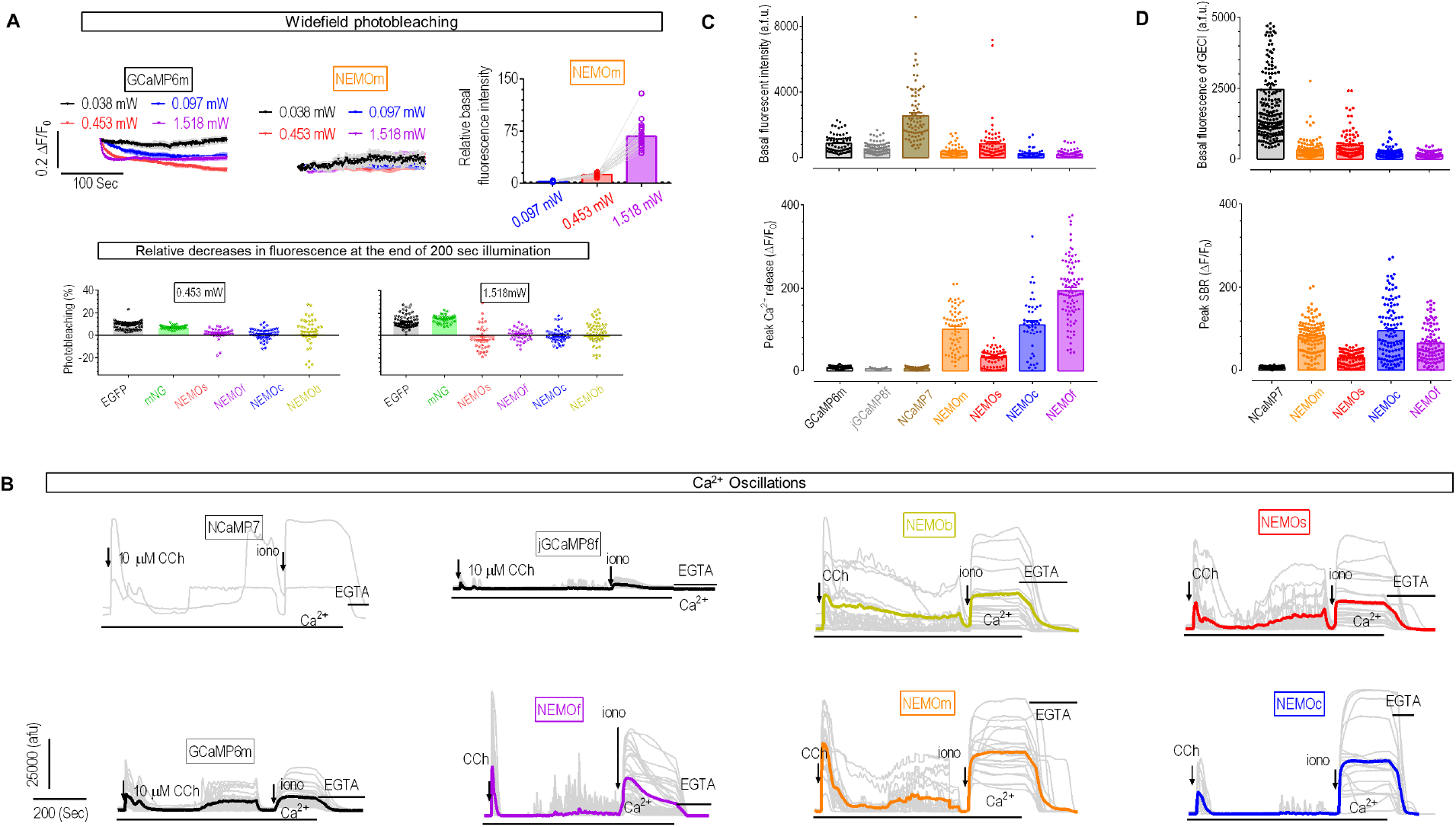
NEMO performance in HEK293 cells. (A) Widefield photobleaching curves. GFP excitation light (470±11nm) was used. Top, two panels on the left, GCaMP6m and NEMOm signals; right, basal fluorescence intensity relative to those excited with 0.038 mW light; Bottom panels, statistics showing the relative reduction in fluorescence at the end of 200 sec illumination with dim (left) or strong (right) light. Top right (n=27 cells); Bottom left (mNG, n=42 cells; EGFP, n=76 cells; NEMOs, n=38 cells; NEMOf, n=37 cells; NEMOc, n=50 cells; NEMOb, n=47 cells); Bottom right (mNG, n=46 cells; EGFP, n=72 cells; NEMOs, n=35 cells; NEMOf, n=42 cells; NEMOc, n=47 cells; NEMOb, n=56 cells). (B) Typical fluorescence oscillations of GECI variants induced by CCh (10 µM). Recordings from the same cells (as presented in Fig. 2A) were shown. Grey lines, responses of individual cells; thick lines, mean responses. (C) Statistics of results shown in panel B and Fig. 2A. Left, the basal fluorescence; right, peak of the first Ca^2+^ release. (GCaMP6m, n=73 cells; NEMOs, n=71 cells; NEMOm, n=63 cells; NEMOc, n=46 cells; NCaMP7, n=77 cells; jGCaMP8f, n=93 cells; NEMOf, n=93 cells). (D) Statistics of results shown in Fig. 2B. Left, mean basal fluorescence; right, the peak SOCE response. n=3 independent biological replicates. (NCaMP7, n=192 cells; NEMOc, n=110 cells; NEMOm, n=117 cells; NEMOf, n=98 cells; NEMOs, n=99 cells). Data from (A), (C) and (D) were from 3 independent biological replicates, and shown as mean ±s.e.m..

**Extended Data Figure 6.**
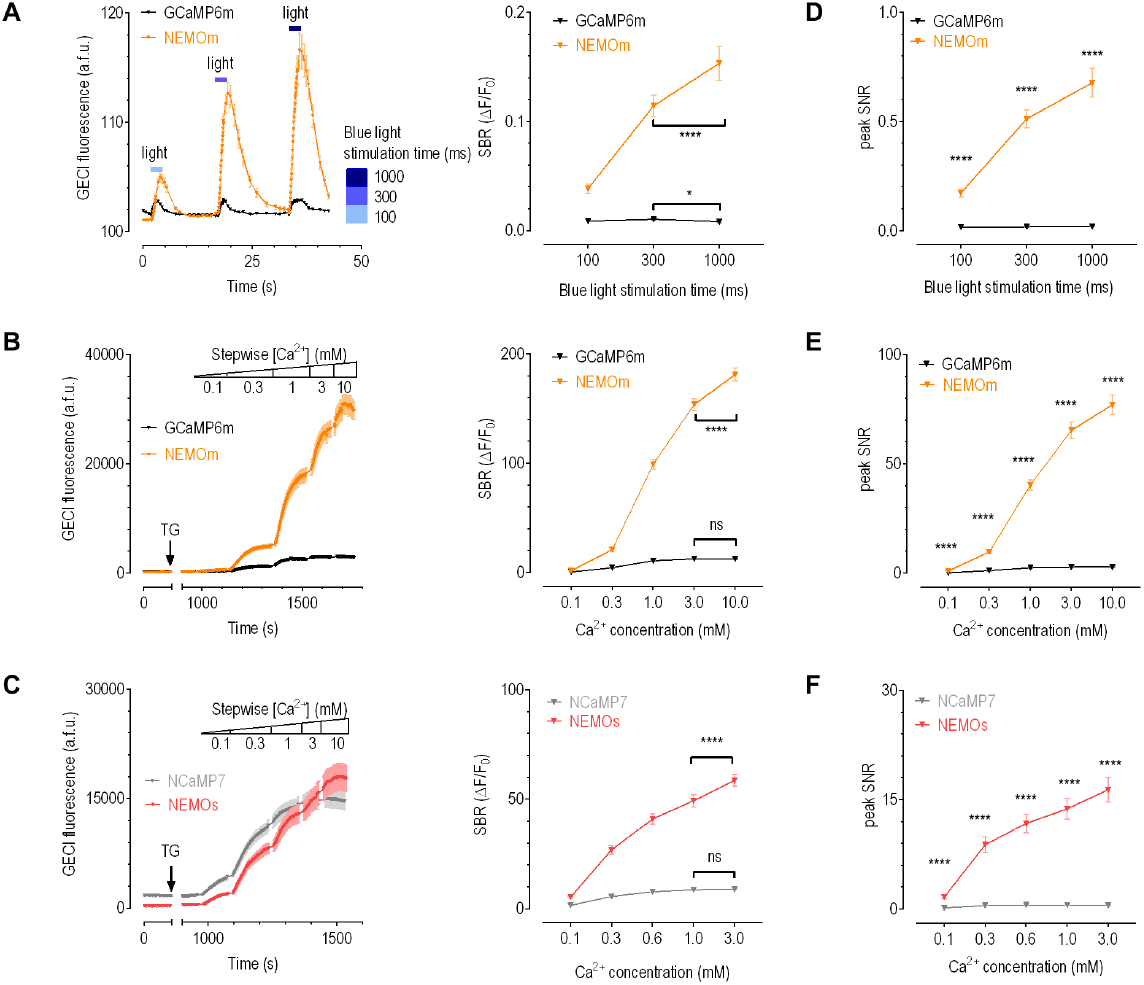
NEMO performance in resolving Ca^2+^ signals that span a wide range of magnitude. (A) Dim blue light stimulation induced GECI responses in HeLa cells co-expressing Opto-CRAC and GCaMP6m or NEMOs. Left, typical traces; right, statistics (NEMOm, 1000 ms versus 300 ms, ****, p<0.0001; GCaMP6m, 1000 ms versus 300 ms, p=0.0178). To avoid direct activation with 488-nm light, minimal (1%) excitation input at 488 nm was used. Three 15-cycles of 2% 473-nm stimulation light with different exposure time were sequentially applied to photo-induce more Ca^2+^ influxes. 100 ms for the first one, 300 ms for the second, and 10000 ms for the third cycle of photo-stimulation. (B-C) SOCE indicated by GECIs in HEK293 cells bathed in stepwise external Ca^2+^. (B, GCaMP6m vs NEMOm; C, NCaMP7 vs NEMOs). Left, typical traces; right, statistics. (For the left panel, NEMOm, 10 mM versus 3 mM, p<0.0001; GCaMP6m, 10 mM versus 3 mM, p=0.7917. For the right panel, NEMOs, 3 mM versus 1 mM, ****, p<0.0001; NCaMP7, 3 mM versus 1 mM, p=0.0681.) (D-F) Statistics of panels A-C showing SNR of GECIs. ****, p< 0.0001. (A-C) Paired Student’s *t*-test, two-tailed. (D-F) Unpaired Student’s *t*-test, two-tailed. n=3 independent biological replicates. All data in this figure were shown as mean ± s.e.m..

**Extended Data Figure 7.**
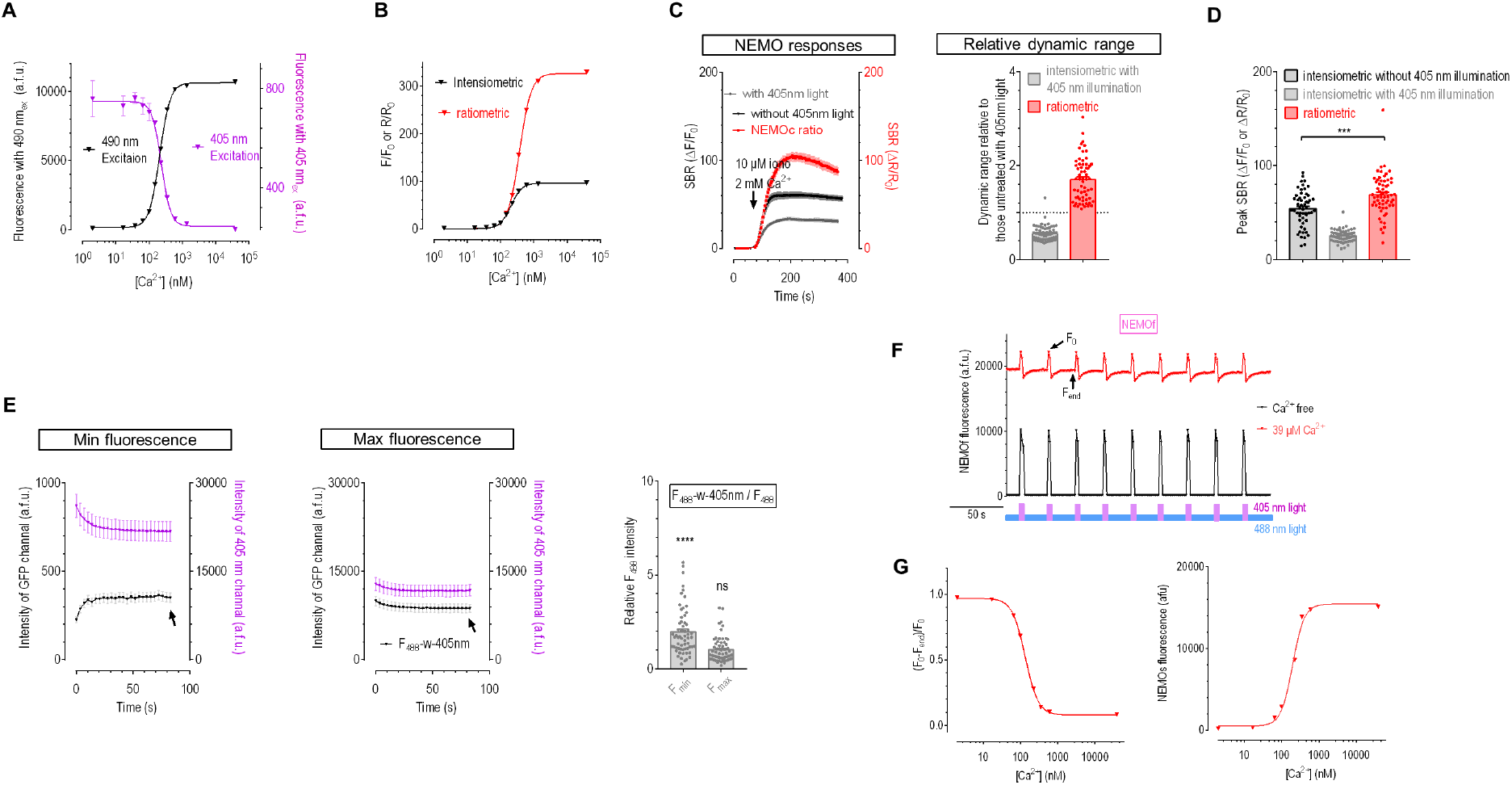
Ratiometric or photochromic responses of NEMO sensors. (A-B) In vitro Ca^2+^ titration curves. (A) 405 nm (F_405_) or 490 nm (F_490_) excited fluorescence responses. n=3 independent measurements. Error bars correspond to s.e.m.. (B) Comparison of instensiometric or ratiometric (R=(F_490_/F_405_) responses of NEMOs. n=3 independent measurements. (C-E) Evaluation of ratiometric responses of NEMOs expressed in HEK cells. 3 independent biological repeats. (C) Maximal NEMOs responses with or without 405 nm illumination. Left, typical traces; right, statistics. (D) Statistics of results in Fig. 2D. (***, p=0.0002, unpaired Student’s *t*-test, two-tailed) (Intensiometric without 405 nm illumination, n=54 cells; intensiometric with 405 nm illumination and ratiometric, n=61 cells). (E) Effect of 405 nm light illumination on F_405_ or F_488_. Left, representative NEMOs responses when cytosolic Ca^2+^ were mostly exhausted by 10 min bath in imaging solution containing 10 μM ionomycin and 1 mM EGTA. Middle, typical traces showing NEMOc responses bathed in 10 μM ionomycin and 2 mM Ca^2+^. Right, statistics. F_488_-w-405nm, NEMOs signal excited with 488nm laser that were exposed to 405 nm (F_min 405_ versus F_488_ and F_max 405_ versus F_488_, ****, p <0.0001 and p=0.88, unpaired Student’s t-test, two-tailed; n=61 cells). (C, E; n=61 cells) (F-G) *In vitro* Ca^2+^ titration of NEMOf with intermittent photochromism-enabled absolute quantification (iPEAQ) method. n=3 independent measurements. (F) Typical traces showing the responses of 488nm-excited NEMOf to repeated illumination by 405 nm light. F_0_ and F_end_, indicated by arrows, peak and minimum fluorescence intensities used to calculate values of the photochromism contrast (F_0_ - F_end_) / F_end_, or (ΔF / F_0_)_hv_. (G) Ca^2+^ titration curves. Left panel, the photochromism contrast ((ΔF / F_0_)_hv_ - Ca^2+^ titration curve that enabled conversion of measured basal photochromism contrast to basal Ca^2+^ concentration of a NEMOs-expressing cell; Right panel, fluorescence - Ca^2+^ titration curve that enabled the subsequent determination of changes in absolute Ca^2+^ concentrations with the calculated basal Ca^2+^ level and recorded NEMOs fluorescence response curve. Except for (F), all other panels in this figure were shown as mean ± s.e.m..

**Extended Data Figure 8.**
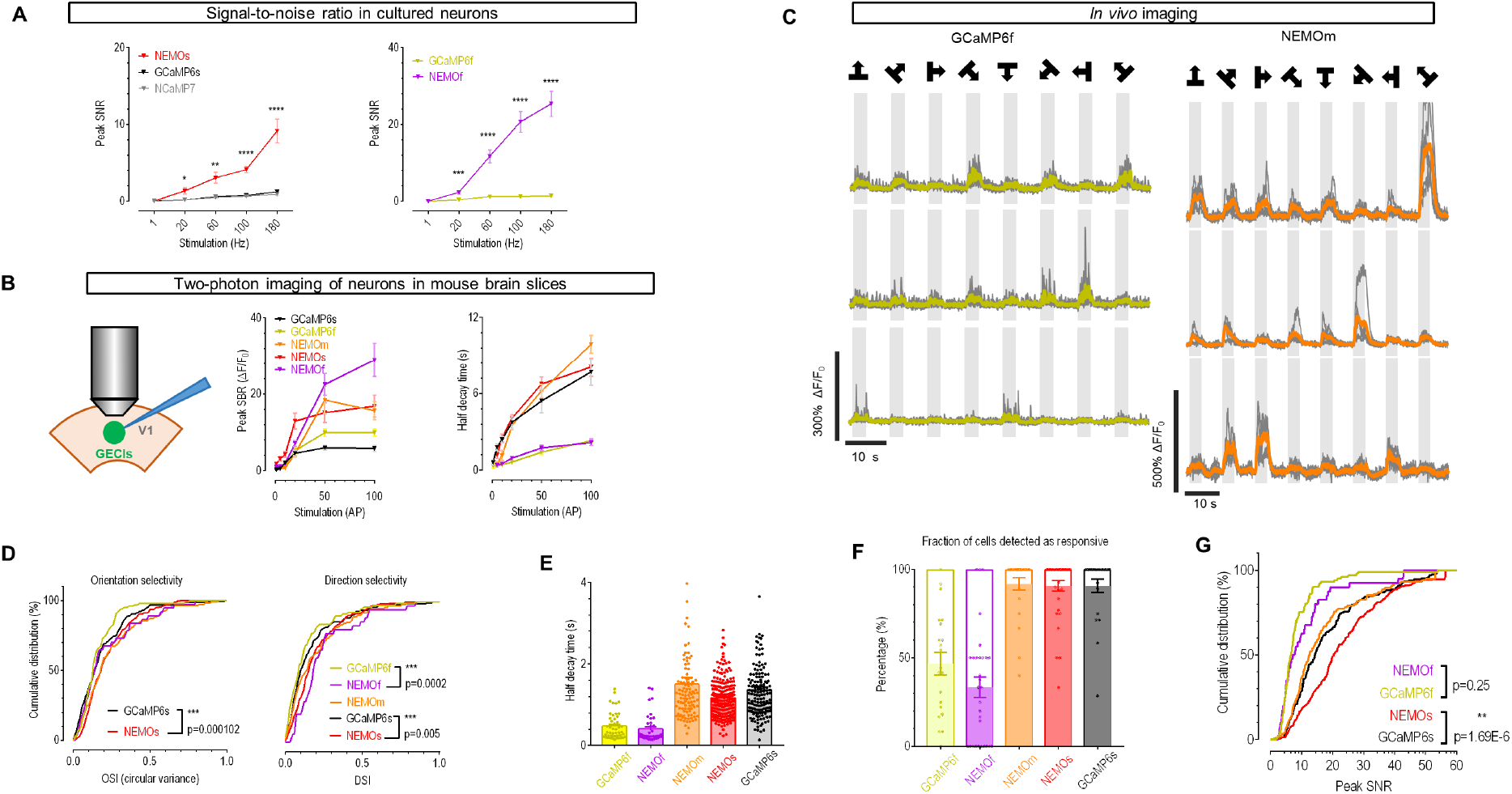
Performance of NEMO sensors in rodent neurons. (A) Signal-to-noise ratio (SNR) of GECIs in cultured neurons (unpaired Student’s *t*-test, two-tailed; Left panel, NEMOm versus NCaMP7 or GCaMP6m from 1 to 180 Hz, p>0.8; **, p=0.029; **, p<0.0015; ****, p<0.0005; ****, p<0.0002; ^****^, p<0.0001; right panel, for 1 and 20Hz, p= 0.7784; for the other frequencies, p<0.0001) (For stimulation at varied frequencies, NCaMP7, n=12, 12, 12, 14, 12 cells. The “n” values of other sensors was equal to Fig.3. Each GECI measurement set was analyzed from multiple dendrites of neurons in three different primary hippocampal neuron cultures. (B) Fluorescence responses of NEMO variants in neurons of acute mouse brain slices. Left, cartoon showing the setup of two-photon fluorescence imaging under a whole-cell patch clamp configuration. Middle, statistics showing the peak SBR-frequency of action potentials (AP) (GCaMP6f, n=57 cells from 3 mice; GCaMP6s, n=51 cells from 2 mice; NEMOs, n=30 cells from 4 mice; NEMOm, n=45 cells from 3 mice; NEMOf, n=54 cells, from 3 mice); right, half decay time of NEMO variants under different intensities of stimuli (For stimulation from 5-100 AP, GCaMP6f, n=12,14,18,18,18 cells from 3 mice; GCaMP6s, n=17 cells from 2 mice; NEMOm, n=2,4,14,16,16 cells from 3 mice; NEMOs, n=8,9,10,10,9 cells from 4 mice; NEMOf, n=5,9,15,18,17 cells from 3 mice; for stimulation at 1 AP, GCaMP6s, n=10 cells from 2 mice; GCaMP6f, n=3 cells from 3 mice). (C-F) *In vivo* two-photon imaging of visual cortex neurons in response to drift gratings. (C) Typical responses; (D-F) Statistics of results shown in Fig. 4A-4C; (D) Cumulative distribution of orientation (left, ^****^, p=0.000102) or direction (right, for NEMOf versus GCaMP6f and NEMOs versus GCaMP6s, ^****^, p = 0.0002; ^***^, p=0.005) selectivity (Kolmogorov–Smirnov test, two-tailed). (E) Statistics of half decay time (GCaMP6f, n=46 cells from 2 mice; GCaMP6s, n=157 cells from 3 mice; NEMOs, n=223 cells from 4 mice; NEMOm, n=105 cells from 3 mice; NEMOf, n=40 cells from 3 mice). (F) Statistics showing fraction of responsive cells (GCaMP6f, n=21 cells from 2 mice; GCaMP6s, n=23 cells from 3 mice; NEMOs, n=35 cells from 4 mice; NEMOm, n=24 cells, from 3 mice; NEMOf, n=30 cells from 3 mice). (G) Cumulative distribution of SNR (NEMOf versus GCaMP6f and NEMOs versus GCaMP6s, p = 0.25; ^****^, p=1.69E-6, Kolmogorov-Smirnov test, two-tailed). At least n=3 independent biological replicates. Data in A, B, E and F panels were shown as mean ± s.e.m..

**Extended Data Figure 9.**
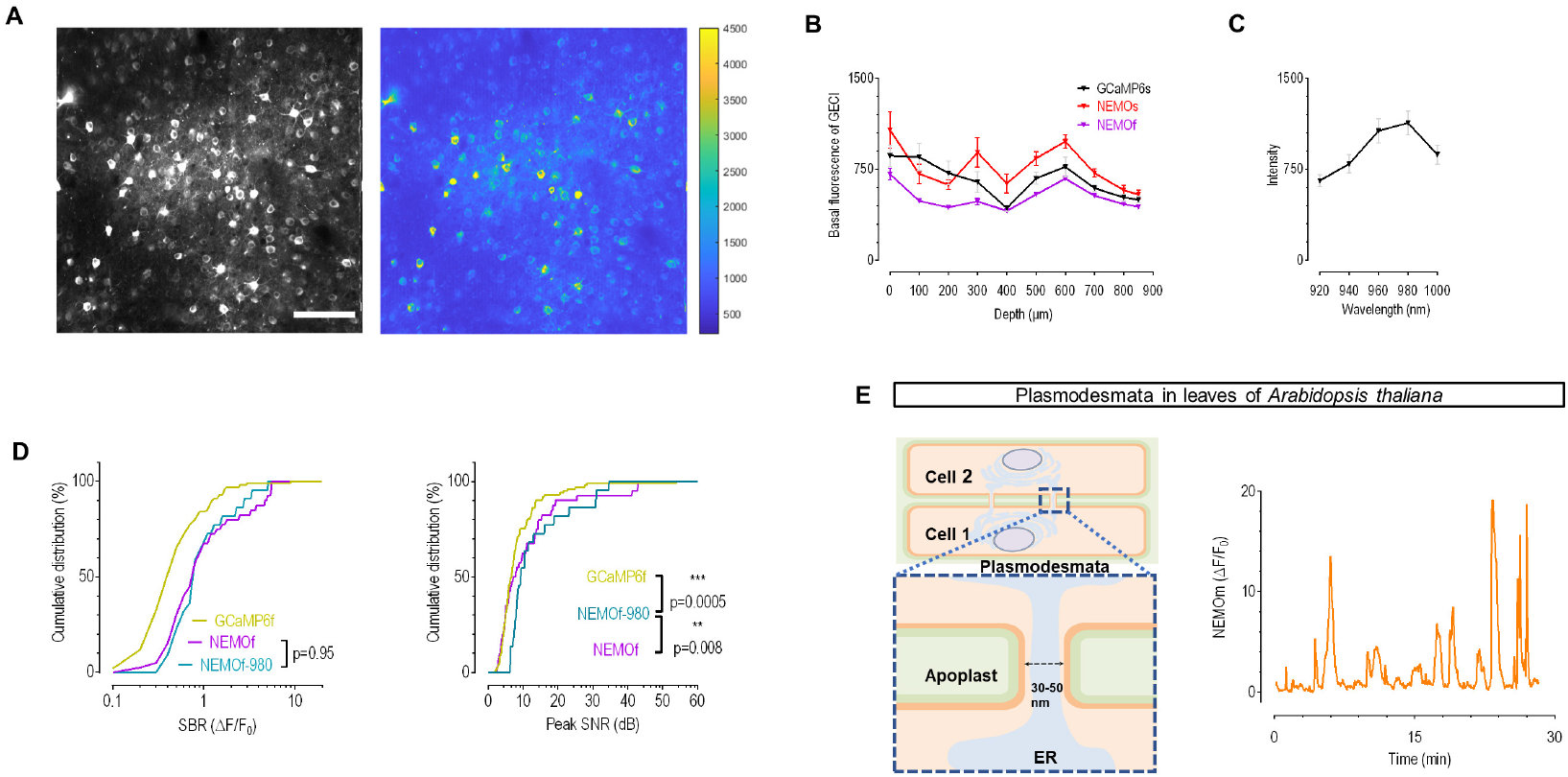
Performance of NEMO in visual cortex neurons from mouse and leaves of *Arabidopsis thaliana*. (A) Typical two-photon images of NEMOs-expressing neurons obtained from the visual cortex of mice. 920-nm light was used to excite NEMOs. (B) Comparison of the basal fluorescence of NEMOs, NEMOf and GCaMP6s under different depth beneath the surface of mouse brain. 920-nm light was used (n=8 ROIs from 2 mice for per sensor). (C) More than two months after infection, the basal fluorescence of NEMOm of the same set of cells excited by light at different wavelengths was quantified (n=20 cells). Data in (B) and (C) were shown as mean ± s.e.m.. (D) Cumulative distribution of SBR (left) or SNR (right) of NEMOf excited at 920-nm (n=112 cells) or 980 nm (n=104 cells). GCaMP6f and NEMOf data were replotted from Fig. S7G. (Left, NEMOf vs NEMOf-980, p=0.95; Right, GCaMP6f vs NEMOf-980, ^***^, p=0.0005; NEMOf vs NEMOf-980, ^**^, p=0.008; Kolmogorov-Smirnov test, two-tailed). E) Ca^2+^ oscillations near plasmodesmata in the leaves of *Arabidopsis thaliana* when excited at 488 nm. NEMOm was tagged with PDLP1, a plasmodesmata marker. Scale bar, 5 μm. Left, cartoon illustration showing the structure of plasmodesmata; right, typical traces. At least n=3 independent biological replicates.

---

**Supplementary Table 1.**
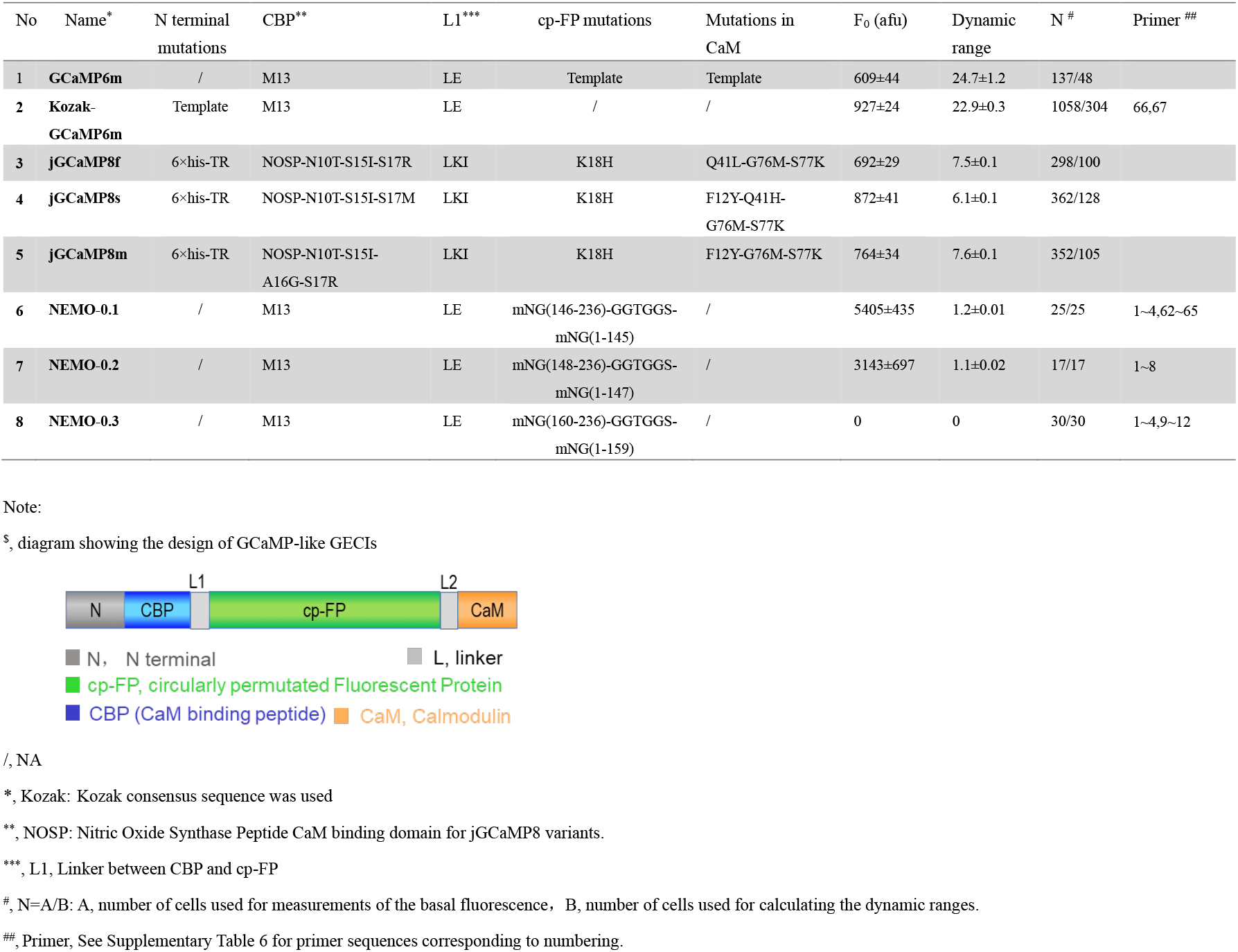
*In cellulo* screening results of constructs with a GCaMP-like design^$^

**Supplementary Table 2.**
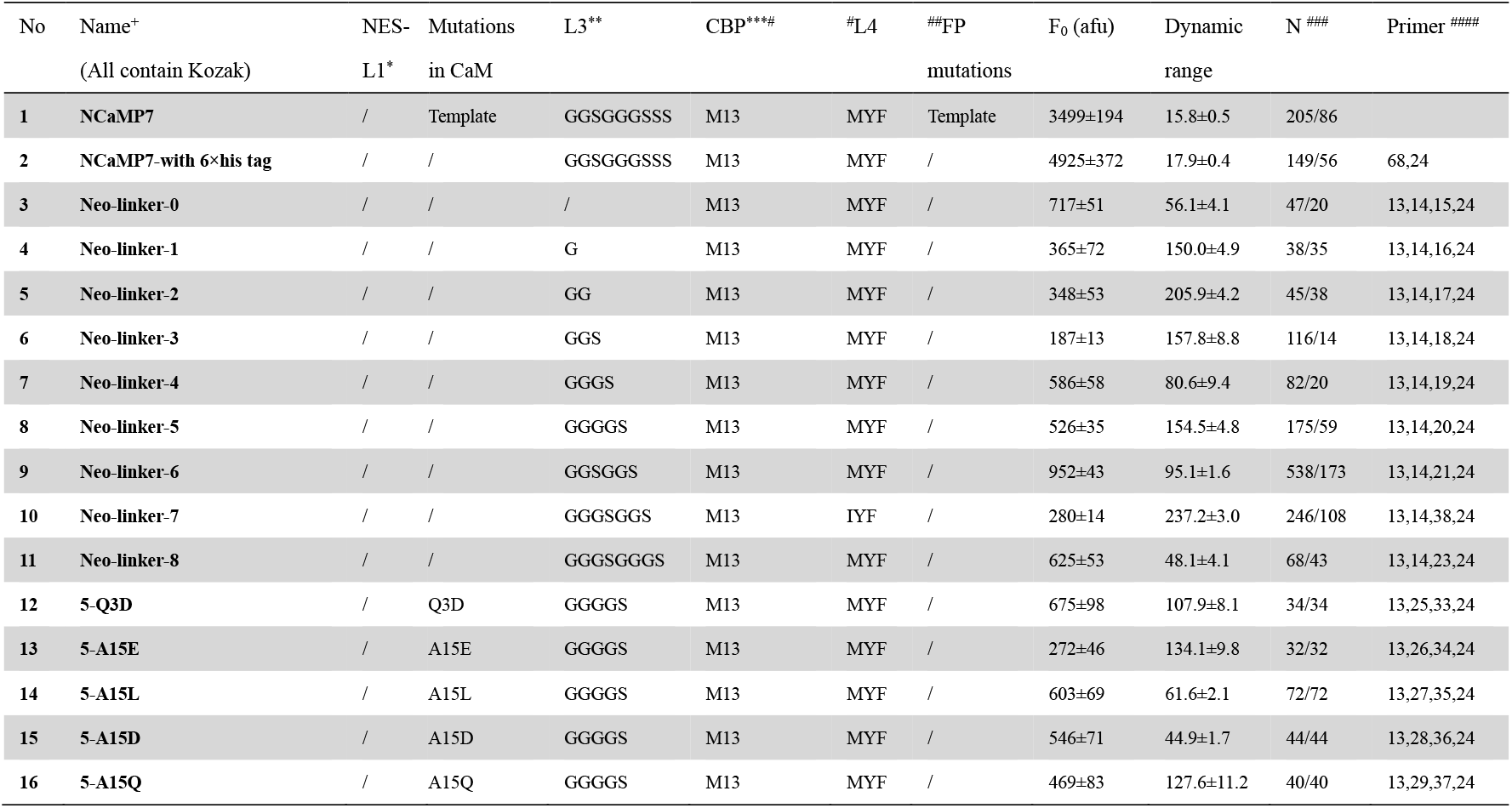
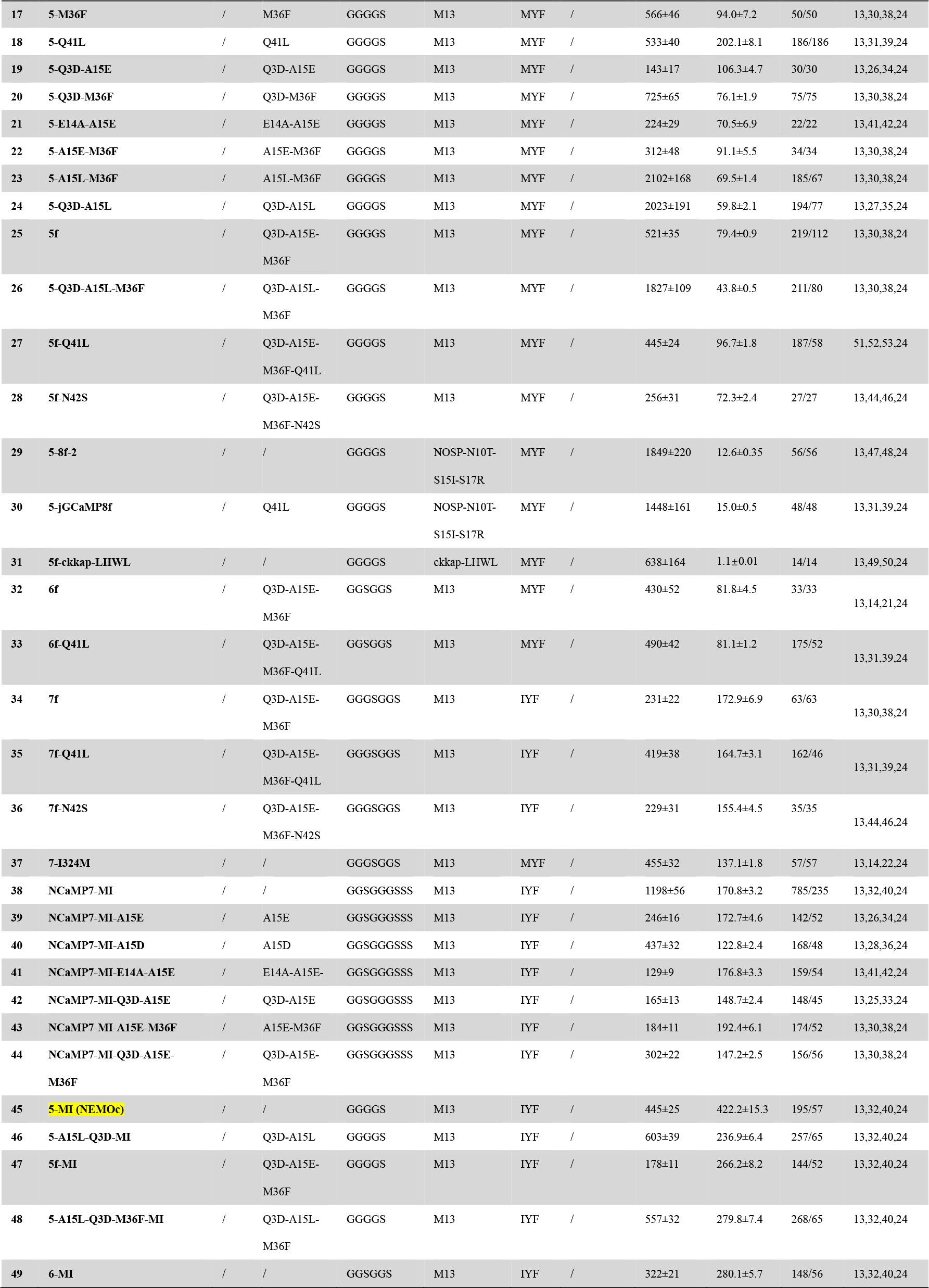
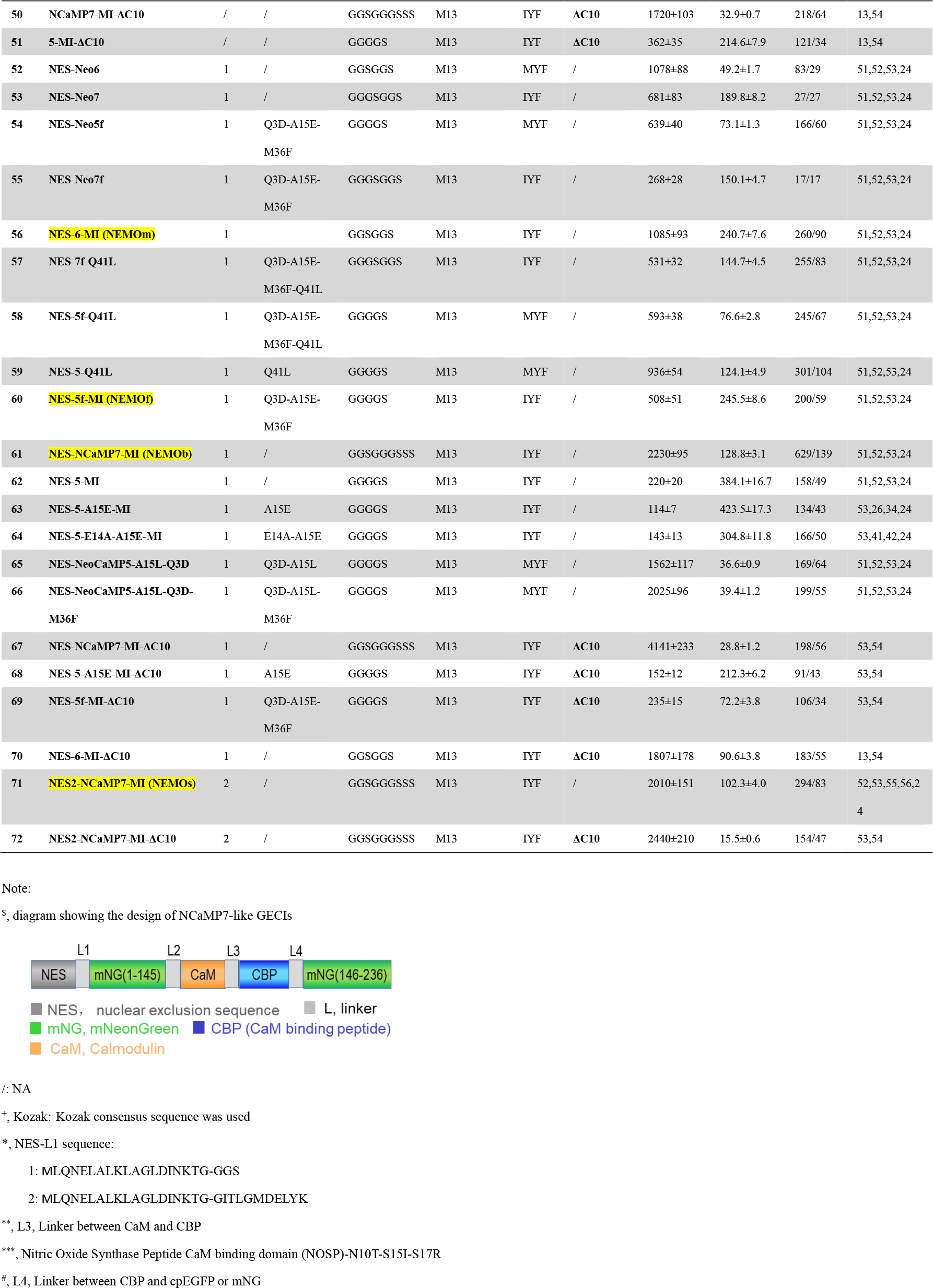
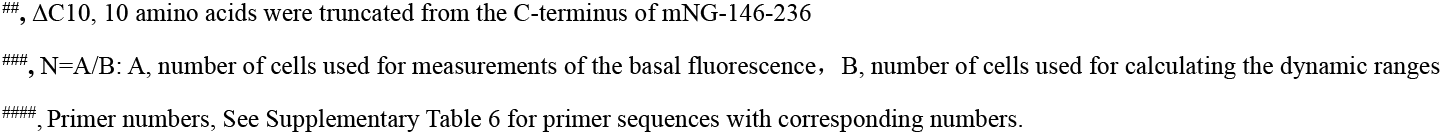
*In cellulo* screening results of NEMO constructs with a NCaMP7-like design^$^

**Supplementary Table 3.** *In vitro* Ca^2+^ titration results and kinetics of NEMO variants

| Sensor | Mg<sup>2+</sup> | K<sub>d</sub> (nM) | Hill number | F<sub>min</sub><br>(10 mM EGTA) | F<sub>rest</sub><br>(100 nM Ca<sup>2+</sup>) | F<sub>max</sub><br>(38 μM Ca<sup>2+</sup>) | Dynamic range<br>(F<sub>max</sub>/F<sub>min</sub>) | K<sub>off</sub> (s<sup>-1</sup>) | K<sub>on</sub><sup>1</sup><br>(*10<sup>6</sup>, M<sup>-1</sup>s<sup>-1</sup>) |
| --- | --- | --- | --- | --- | --- | --- | --- | --- | --- |
| GCaMP6m | - | 174.3±6.1 | 2.5±0.2 | 155±6 | 700±49 | 4023±93 | 26.7±0.9 | 2.48±0.02 | 14.20 |
|  | + | 206.4±5 | 2±0.1 | 176±4 | 680±34 | 4133±85 | 23.6±0.6 | 2.52±0.02 | 12.21 |
| GCaMP6f | - | 354.4±1.5 | 2±0.2 | 228±4 | 515.8±8.9 | 8019±252 | 35.2±1.3 | 3.72±0.004 | 10.49 |
|  | + | 568.8±9.4 | 1.7±0.0 | 268±3 | 477.8±4.4 | 8250±212 | 30.7±0.6 | 4.35±0.03 | 7.65 |
| NCaMP7 | - | 97.9±2 | 2.5±0.1 | 406±29 | 16870±1964 | 32753±2394 | 80±3.2 | 3.29±0.01 | 33.56 |
|  | + | 98.3±1.5 | 2.4±0.1 | 677±45 | 16216±1325 | 30628±1927 | 45.5±0.9 | 2.48±0.01 | 25.23 |
| NEMOc | - | 492.8±15.9 | 3.4±0.3 | 98±4 | 152±4 | 15875±236 | 165.3±7.7 | 4.59±0.03 | 9.31 |
|  | + | 557.3±15.5 | 3.8±0.4 | 89±7 | 137±6 | 15267±578 | 182.2±12.6 | 3.16±0.02 | 5.67 |
| NEMOm | - | 198.9±4.8 | 3.3±0.2 | 123±5 | 1735±150 | 19396±725 | 159.2±4.3 | 2.22±0.01 | 11.17 |
|  | + | 248.4±3.7 | 3.3±0.1 | 134±6 | 1065±21 | 19706±520 | 149.1±3.5 | 1.46±0.01 | 5.88 |
| NEMOf | - | 440.1±19.4 | 2.9±0.3 | 118±1 | 205±13 | 16705±411 | 141.5±3.8 | 6.73±0.04 | 15.30 |
|  | + | 528.4±17 | 3.2±0.3 | 118±2 | 304±53 | 15085±265 | 127.9±2.3 | 5.57±0.04 | 10.54 |
| NEMOs | - | 129.1±3.1 | 3.1±0.2 | 65±3 | 2572±254 | 8351±194 | 131.3±3.4 | 2.26±0.01 | 17.48 |
|  | + | 155.8±3.8 | 2.7±0.2 | 87±5 | 1890±115 | 8489±311 | 99.7±2.9 | 1.46±0.01 | 9.37 |
| NEMOb | - | 198.7±2.6 | 2.5±0.1 | 340±1 | 3880±89 | 27829±713 | 81.9±2.1 | 1.96±0.003 | 9.85 |
|  | + | 141±1.3 | 2.6±0.1 | 414±4 | 8019±54 | 27857±408 | 67.4±1.3 | 1.56±0.01 | 11.06 |

Note: <sup>1</sup> K<sub>on</sub> was calculated from K<sub>off</sub> and K<sub>d</sub> with this equation: K<sub>on</sub> = K<sub>off</sub>/K<sub>d</sub>

**Supplementary Table 4.** *In vitro* biophysical properties of NEMO sensors

| Sensor | *1Mg<sup>2+</sup> | Ca<sup>2+</sup> | ρ<sup>*2</sup> |  | ε<sub>max</sub> (mM<sup>-1</sup>*cm<sup>-1</sup>) |  | Φ |  | F<sub>1</sub><sup>*3</sup> (mM<sup>-1</sup>*cm<sup>-1</sup>) |  | pKa |
| --- | --- | --- | --- | --- | --- | --- | --- | --- | --- | --- | --- |
|  |  |  | Anionic | Neutral | Anionic | Neutral | Anionic | Neutral | Anionic | Neutral |  |
| GCaMP6m | - | + | 0.47±0.00 | 0.53±0.00 | 82.18±1.48 | 40.60±0.65 | 0.65 | 0.14 | 25.10±0.45 | 3.01±0.05 | 7.18±0.09 |
|  |  | - | 0.05±0.05 | 0.95±0.01 | 60.89±12.06 | 40.10±1.35 | 0.38 | 0.09 | 1.15±0.05 | 3.42±0.13 | / |
|  | + | + | 0.46±0.02 | 0.54±0.02 | 77.99±2.95 | 38.16±4.43 | 0.66 | 0.14 | 23.66±1.22 | 2.87±0.26 | 7.27±0.17 |
|  |  | - | 0.04±0.01 | 0.96±0.01 | 85.96±13.50 | 35.89±1.56 | 0.38 | 0.09 | 1.11±0.13 | 3.11±0.11 | / |
| NCaMP7 | - | + | 0.84±0.02 | 0.16±0.02 | 116.40±2.20 | 42.22±4.71 | 0.75 | 0.16 | 73.67±2.70 | 1.03±0.05 | 6.07±0.62 |
|  |  | - | 0.21±0.01 | 0.79±0.01 | 14.99±0.33 | 51.92±2.50 | 0.35 | 0.08 | 1.12±0.07 | 3.26±0.10 | / |
|  | + | + | 0.85±0.02 | 0.15±0.02 | 107.10±0.91 | 38.60±3.12 | 0.72 | 0.18 | 65.63±1.73 | 1.02±0.04 | 4.93±0.33 |
|  |  | - | 0.22±0.01 | 0.78±0.01 | 15.29±1.26 | 52.16±1.95 | 0.40 | 0.08 | 1.34±0.06 | 3.26±0.16 | / |
| NEMOc | - | + | 0.85±0.02 | 0.15±0.02 | 113.20±3.44 | 40.31±1.14 | 0.67 | 0.18 | 64.54±3.02 | 1.08±0.08 | 5.07±0.33 |
|  |  | - | 0.20±0.02 | 0.80±0.02 | 7.35±1.13 | 49.56±0.91 | 0.29 | 0.06 | 0.43±0.06 | 2.37±0.01 | / |
|  | + | + | 0.86±0.01 | 0.14±0.01 | 111.40±0.91 | 41.81±1.67 | 0.67 | 0.17 | 64.26±2.67 | 0.99±0.05 | 4.97±0.43 |
|  |  | - | 0.19±0.01 | 0.81±0.01 | 7.68±0.25 | 50.07±1.59 | 0.15 | 0.07 | 0.22±0.01 | 2.84±0.06 | / |

Note: \*1 The effect of Mg<sup>2+</sup> was tested because the free cytosolic Mg<sup>2+</sup> concentration is in the range of mM and the presence of Mg<sup>2+</sup> might affect the dynamic range of NCaMP7 (Table 1 in Subach et al, 2020), the parental template of NEMO sensors. \*2 ρ is the relative concentration of chromophore; \*3 F<sub>1</sub> is one-photon brightness defined as the product of ρ, ε and Φ.

**Supplementary Table 5.** Two-photon biophysical properties of NEMO sensors *in vitro*.

| Sensor | Mg<sup>2+</sup> | Ca<sup>2+</sup> | σ<sub>2</sub><sup>*1</sup> (GM) |  | F<sub>2</sub><sup>*2</sup> (GM) |  |
| --- | --- | --- | --- | --- | --- | --- |
|  |  |  | Anionic | Neutral | Anionic | Neutral |
| GCaMP6m | + | + | 18.18±1.2 | 24.41±1.55 | 5.52±0.37 | 1.85±0.12 |
| NCaMP7 | + | + | 42.37±4.13 | 16.9±3.69 | 25.93±2.53 | 0.46±0.10 |
| NEMOc | + | + | 51.94±6.07 | 21.59±5.01 | 29.93±3.50 | 0.51±0.12 |

Note: \*1 σ<sub>2</sub> is two-photon cross-sections. \*2 F<sub>2</sub> is two-photon brightness defined as the product of ρ, Φ and σ<sub>2</sub>; ρ, Φ are from Supplementary table 4.

**Supplementary Table 6.**
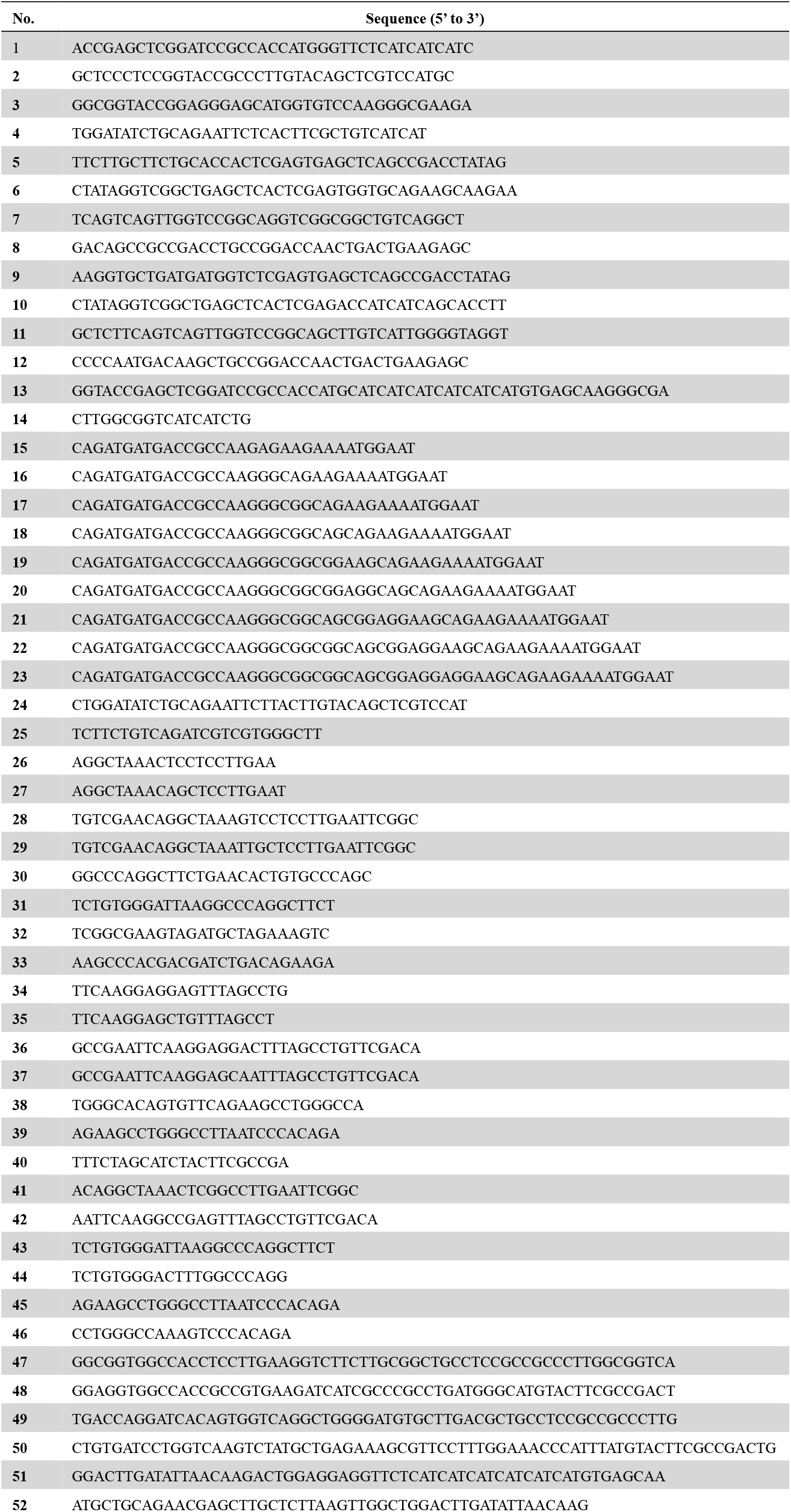
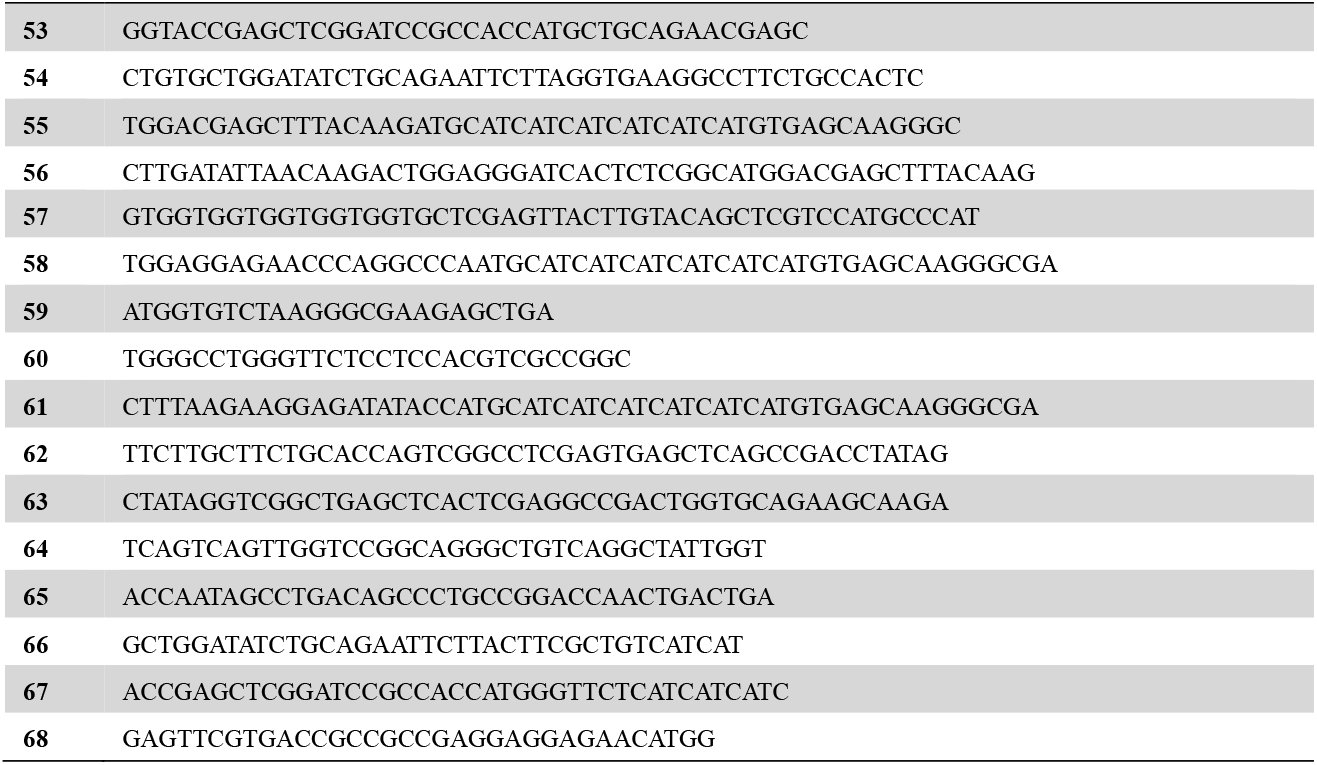
Sequences of primers used to generate GECI constructs listed in Supplementary Table 1&2.

## Movie legends

**Supplementary Video 1. Typical Ca^2+^ release and SOCE response shown by GCaMP6m (left) or NEMOc (right) in HEK293 cells**. Endoplasmic reticulum Ca^2+^ store was depleted using 2.5 μM ionomycin and 1 μM thapsigargin. 100 mM Ca^2+^ was applied to achieve maximal SOCE response.

**Supplementary Video 2. Typical Ca^2+^ oscillations reported by GCaMP6m (left) or NEMOf (right) in HEK293 cells**. Carbachol (CCh, 10 μM) was used to trigger Ca^2+^ oscillation.

**Supplementary Video 3. Representative light induced-Ca^2+^ influxes monitored by GCaMP6m (left) or NEMOm (right) in HeLa cells co-expressing Opto-CRAC, which enables light-inducible activation of endogenous ORAI Ca^2+^ channels**. To avoid activation of Opto-CRAC by 488 nm light used to detect GECI fluorescence, dim light input (1% power of 488-nm laser) and short exposure (10 ms) were used. The cells were sequentially subjected to increased duration of photo-stimulation (100, 300, and 1000 ms) to evoke larger Ca^2+^ influx stepwise.

**Supplementary Video 4. Representative NEMOm-reported Ca^2+^ oscillations adjacent to plasmodesmata in the leaves of *Arabidopsis thaliana***. Scale bar, 5 μm.

## Notes

### Competing Interest Statement

The authors have declared no competing interest.

### Summary of Updates

Following suggestions from reviewers, and editing requirements from Nature Methods journal.

